# Single-cell transcriptome profiling reveals neutrophil heterogeneity and orchestrated maturation during homeostasis and bacterial infection

**DOI:** 10.1101/792200

**Authors:** Xuemei Xie, Qiang Shi, Peng Wu, Xiaoyu Zhang, Hiroto Kambara, Jiayu Su, Hongbo Yu, Shin-Young Park, Rongxia Guo, Qian Ren, Sudong Zhang, Yuanfu Xu, Leslie E. Silberstein, Tao Cheng, Fengxia Ma, Cheng Li, Hongbo R. Luo

**Author notes:** To whom all correspondence should be addressed. Enders Research Building, Room 811 Boston, MA 02115, USA Hongbo R. Luo, (Lead Contact); Cheng Li; Fengxia Ma. These authors contributed equally to this work (Xuemei Xie, Qiang Shi).

## Abstract

The full neutrophil heterogeneity and differentiation landscape remains incompletely characterized. Here we profiled >25,000 differentiating and mature mouse neutrophils using single-cell RNA sequencing to provide a comprehensive transcriptional landscape of neutrophil maturation, function, and fate decision in their steady state and during bacterial infection. Eight neutrophil populations were defined by distinct molecular signatures. The three mature peripheral blood neutrophil subsets arise from distinct maturing bone marrow neutrophil subsets. Driven by both known and uncharacterized transcription factors, neutrophils gradually acquire microbicidal capability as they traverse the transcriptional landscape, representing an evolved mechanism for fine-tuned regulation of an effective but balanced neutrophil response. Bacterial infection reprograms the genetic architecture of neutrophil populations, alters dynamic transition between each subpopulation, and primes neutrophils for augmented functionality without affecting overall heterogeneity. In summary, these data establish a reference model and general framework for studying neutrophil-related disease mechanisms, biomarkers, and therapeutic targets at single-cell resolution.

**Graphical Abstract:** 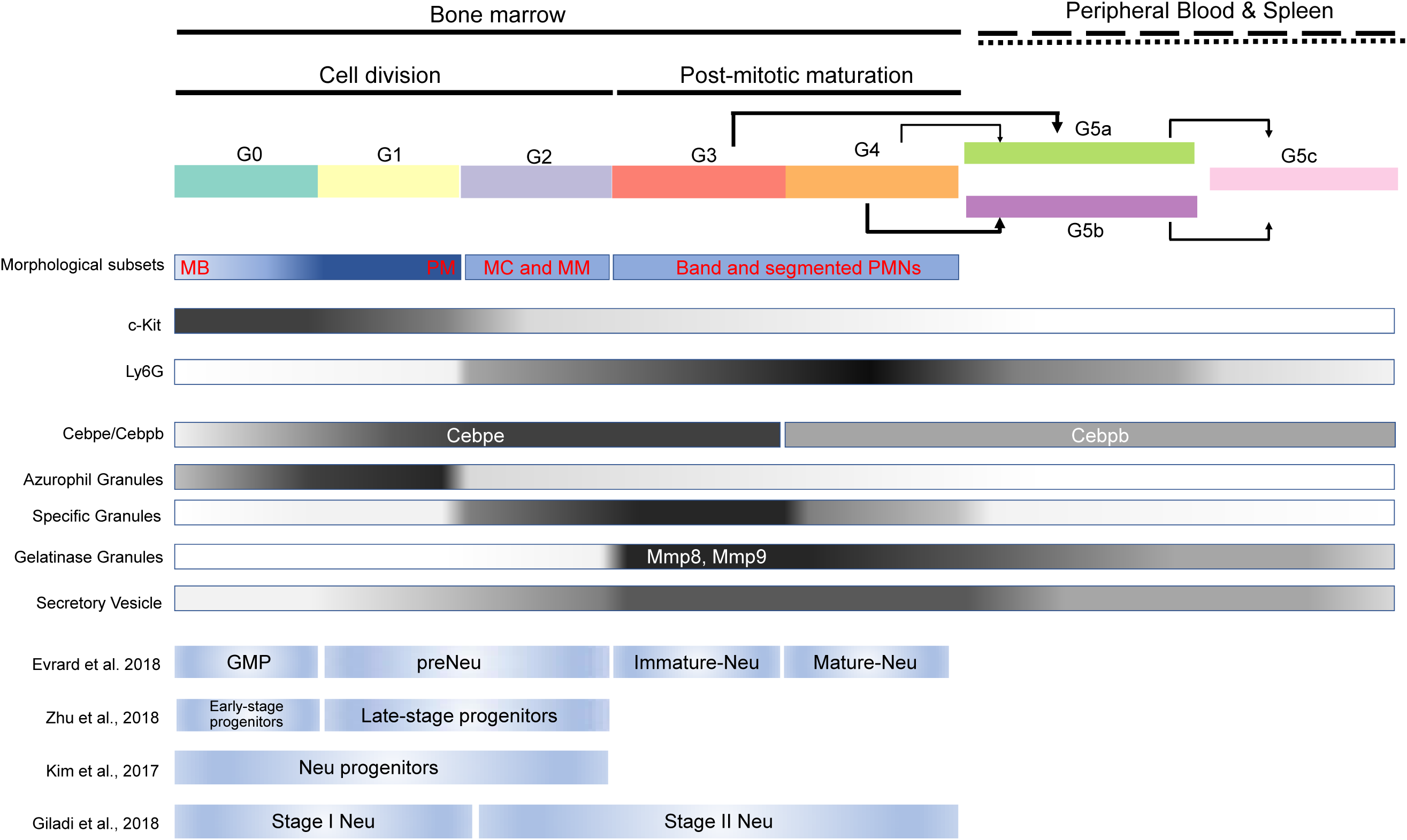

**Highlights:**

- A comprehensive single-cell resolution transcriptional landscape of mouse neutrophil maturation and fate decision under steady-state and bacterial infection conditions.
- The pathogen clearance machinery in neutrophils is continuously and gradually built during neutrophil differentiation, maturation, and aging, driven by both known and uncharacterized transcription factors.
- The three mature neutrophil subsets in peripheral blood, including a novel ISG-expressing subset, are derived from distinct bone marrow neutrophil precursors.
- Bacterial infection reprograms the genetic architecture of neutrophil populations, alters dynamic transition between each subpopulation, and primes neutrophils for augmented functionality without affecting overall neutrophil heterogeneity.
- Bacterial infection-induced emergency granulopoiesis is mediated by augmented proliferation of early stage neutrophil progenitors and accelerated post-mitotic maturation.

## Introduction

Neutrophils migrate from the circulating blood to infected tissues in response to inflammatory stimuli, where they protect the host by phagocytosing, killing, and digesting bacterial and fungal pathogens (Cossio et al., 2019; Darrah and Andrade, 2012; Lee et al., 2003; Ley et al., 2018; Nauseef and Borregaard, 2014; Nicolas-Avila et al., 2017; Segal, 2005). Neutrophil function must be tightly controlled during infection and inflammation: excessive neutrophil accumulation or hyper-responsiveness can be detrimental (Baggiolini, 2001; Castanheira and Kubes, 2019; Davis et al., 2003; Wipke and Allen, 2001), while defects in neutrophil development, trafficking, or function can result in immunological and hematological disorders (Bunting et al., 2002; Burg and Pillinger, 2001; Dinauer, 2019; Fodil et al., 2016; Kruger et al., 2015; Witko-Sarsat et al., 2000).

In adults, neutrophils are mainly produced in the bone marrow (BM) through stepwise progression of myeloid progenitors (Cowland and Borregaard, 2016a; Lawrence et al., 2018; Ward et al., 2000). Neutrophil maturation typically follows five main morphological stages: myeloblasts (MBs), promyelocytes (PMs), myelocytes (MCs), metamyelocytes (MMs), band neutrophils and segmented neutrophils (BC/SCs). Cell division is halted at an early stage of neutrophil differentiation (Klausen et al., 2004; Mora-Jensen et al., 2011), while post-mitotic maturation after the last cell division occurs in the BM, with mature neutrophils released through the venular endothelium into the bloodstream as non-dividing polymorphonuclear neutrophils (PMNs) (Cowland and Borregaard, 2016a).

Neutrophil populations are not, however, homogenous (Adrover et al., 2016; Ley et al., 2018; Ng et al., 2019; Nicolas-Avila et al., 2017; Scapini et al., 2016; Silvestre-Roig et al., 2016; Yvan-Charvet and Ng, 2019): differentiation and maturation produce distinct neutrophils subpopulations which may be pre-programmed with different functions; discrete microenvironments can modify neutrophil function and behavior; and rapid neutrophil aging, their short lifespan, and mechanically-induced cellular responses as they enter and exit capillaries (Doerschuk, 2000; Wang and Doerschuk, 2002) contribute to neutrophil heterogeneity. Neutrophil classification has traditionally relied on morphology, surface marker expression, or gradient separation, which while simple and robust do not capture the full neutrophil compartment repertoire. Some neutrophil subpopulations overlap, making nomenclature confusing and contributing to controversies regarding neutrophil function and ontogeny. In addition, the exact function of some neutrophil subpopulations and the molecular bases of heterogeneity are varied and remain elusive.

Single-cell RNA sequencing (scRNA-seq) is a powerful tool to explore immune cell heterogeneity (Adlung and Amit, 2018; Papalexi and Satija, 2018; Stubbington et al., 2017). Here we adopt an unbiased, systematic approach to dissect mouse neutrophil populations in the bone marrow, peripheral blood, and spleen at single-cell resolution. In doing so, we provide the first comprehensive reference map of differentiating and mature neutrophil transcriptional states in both healthy and *E. coli*-infected hosts. Our analysis addresses critical deficiencies in our understanding of neutrophil biology, and the resource will provide the wider community with new opportunities to further dissect the complex molecular regulation of neutrophil maturation, function, and fate, including for therapeutic impact.

## Results

### Mouse neutrophil atlas in the steady state

To characterize the gene expression programs dictating neutrophil differentiation, maturation and steady-state heterogeneity, we obtained a comprehensive scRNA-seq map of mouse neutrophils under steady-state conditions. Gr1^+^ cells were isolated from the bone marrow (BM-Gr1), peripheral blood (PB-Gr1), and spleen (SP-Gr1) by FACS (**Fig.1a**). To capture the whole spectrum of neutrophil maturation and identify potential neutrophil populations with lower Gr1 antigen (mainly Ly6G) expression, we also included Gr1^low^ and a few Gr1^-^ cells in each sample. In addition, to gain insights into granulopoiesis in its entirety, we included a sample enriched for c-Kit^+^ BM HSPCs mixed with BM-Gr1 cells at a 2:3 ratio to artificially create a BM-cKit/Gr1 population. The sorted high-quality intact single cells (**Fig.S1a-e**) were processed for RNA-seq using the 10x Genomics Chromium platform (**Fig.1a**).

**Figure 1.**
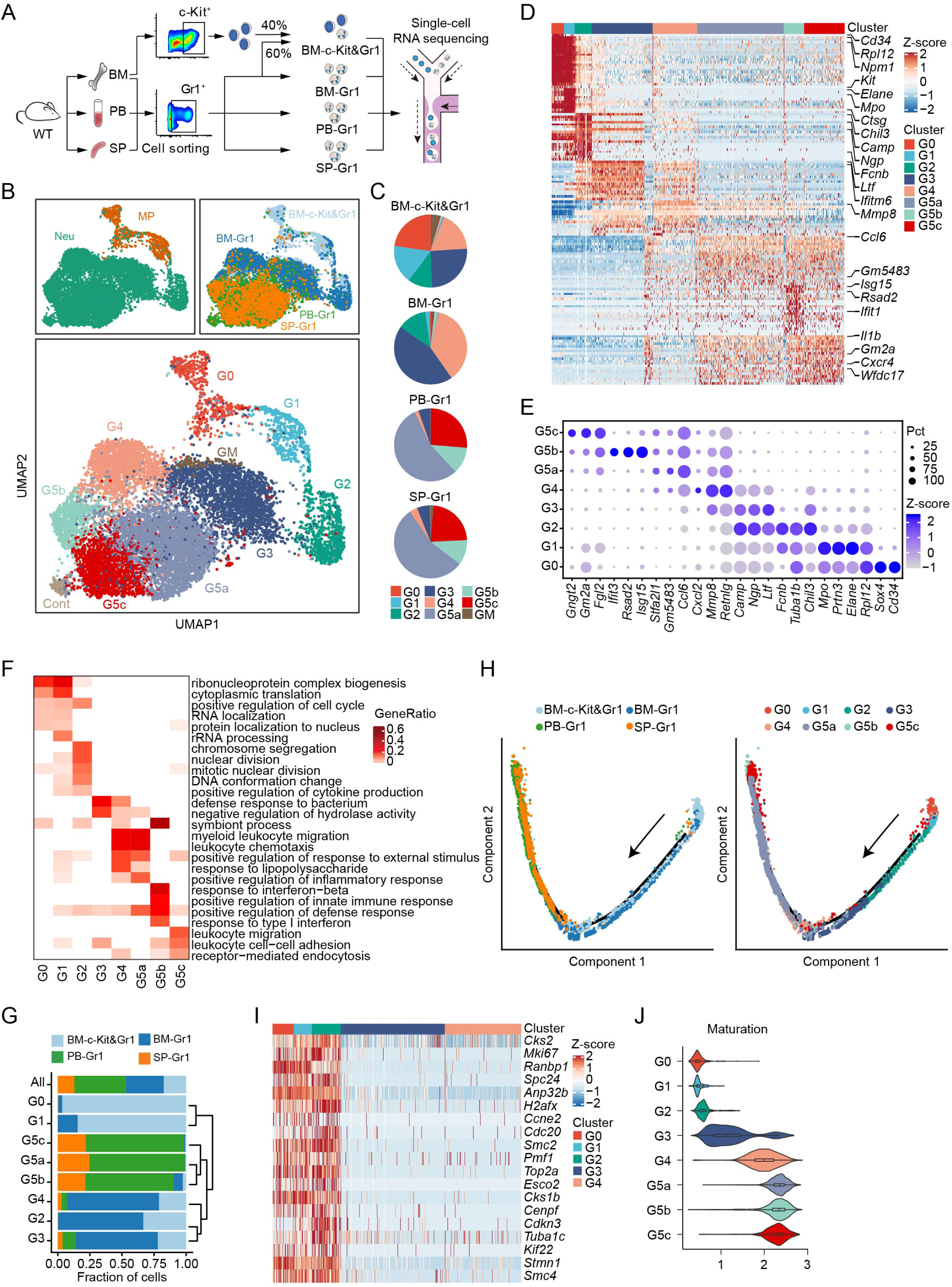
**Single-cell RNA-seq analysis of steady-state bone marrow (BM), peripheral blood (PB), and spleen (SP) neutrophils.** (A) Overview of study design. (B) Uniform manifold approximation and projection (UMAP) of 12,285 neutrophils from BM, PB, and SP colored by cell type, sample origin, and inferred cluster identity, respectively. MP: myeloid progenitors. Cont: the contaminating population mainly consisted of low-quality cells so was discarded from further analysis. GM: low-quality G3-like cells (low UMI count per cell, low UMI count per gene, and high percentage of mitochondrial UMI counts). Similarly, unless otherwise stated, GM were excluded from all other downstream analyses. (C) Proportions of the nine neutrophil clusters in four samples. (D) Heatmap showing row-scaled expression of the 20 highest differentially expressed genes (DEGs, Bonferroni-corrected P-values < 0.05, Student’s t-test) per cluster for all neutrophils except cells from the GM population. (E) Dot plot showing scaled expression of selected signature genes for each cluster colored by average expression of each gene in each cluster scaled across all clusters. Dot size represents the percentage of cells in each cluster with more than one read of the corresponding gene. (F) Gene ontology (GO) analysis of DEGs for each cluster. Selected GO terms with Benjamini-Hochberg-corrected P-values < 0.05 (one-sided Fisher’s exact test) are shown and colored by gene ratio. (G) Unsupervised hierarchical clustering of the eight clusters based on the average gene expression of cells in each cluster. Organ distribution of each cluster is shown on the left. (H) Monocle trajectories of neutrophils colored by sample origin (left) and cluster identity (right). Each dot represents a single cell. Cell orders are inferred from the expression of the most variable genes across all cells. Trajectory directions were determined by biological prior. (I) Heatmap showing row-scaled expression of cell cycle-related genes for G0-G4 neutrophils. (J) Violin plot of maturation scores for each cluster.

To obtain a harmonized atlas, we performed an integrated analysis of cells pooled from different organs (**Fig.S1f**) and projected into two dimensions with Uniform Manifold Approximation and Projection (UMAP) (Becht et al., 2018a) based on their transcriptomic profiles. Unbiased, graph-based clustering identified seven major cell populations (**Fig.S1g-I and Table S2**). Lymphocytes and monocytes from different samples overlapped in the same defined clusters, indicating that there was little batch effect between samples. Since the majority of the cells were Gr1^+^, the largest cell cluster identified by this analysis was the neutrophil population, as intended (**Fig.S1g)**.

To dissect neutrophil heterogeneity, we examined the neutrophil-related populations (MPs and neutrophils). Unsupervised clustering partitioned differentiating and mature neutrophils into eight clusters (G0-4; G5a-c; **Fig.1b**), most containing cells derived from multiple tissues with overlapping UMAP coordinates. G0-4 mainly originated from the BM and represented neutrophils differentiating in the BM, while G5a-c mainly originated from peripheral tissues (**Fig.1c**). There was substantial differential gene expression between the groups (**Fig.1d and Table S3)**. For quality assurance, we merged our raw data with another high-quality published dataset (Giladi et al., 2018) (**Fig.S1j-k**), which agreed with our data well, our data detecting even greater numbers of genes (**Fig.S1k**). We identified 1,098 differentially-expressed genes (DEGs) and 27 signature genes that distinguished each subpopulation (**Fig.1e**). In a gene ontology analysis of DEGs (**Fig.1f**), cell cycle-related genes were as expected highly expressed in earlier phases of neutrophil maturation (G0, G1, and G2). G0 and G1 cells were also enriched for protein synthesis genes related to rRNA processing and protein translation. Interestingly, genes related to cytokine production started to be expressed in dividing G2 cells.

### Neutrophil differentiation and maturation trajectories

Since we intentionally included c-Kit^+^ cells, significant numbers of CD34^+^ G0 cells were detected. According to known gene signatures (Karamitros et al., 2018; Nestorowa et al., 2016; Paul et al., 2015; Velten et al., 2017), we concluded that the G0 population mainly consisted of granulocyte-macrophage progenitors (GMPs) expressing typical genes such as *Cd117* (*Kit*), *Cd34*, and *Sox4* and neutrophil primary granule genes such as *Mpo*, *Elane*, and *Prtn3* (**Fig.1e**). To investigate the interrelationship between neutrophil subpopulations, we conducted hierarchical clustering (**Fig.1g)**. Consistent with UMAP clustering, neutrophils in the PB (G5a, G5b, and G5c) were closely associated but more remote from BM cells (G1-4). Using Monocle (Qiu et al., 2017a) to place differentiating neutrophil populations along possible granulopoiesis trajectories in pseudo-time (**Fig.1h**), neutrophil differentiation and maturation occurred on a tightly organized trajectory, starting from G1 cells in the BM and ending with G5 cells in the PB and the spleen. G1 to G2 cells underwent active proliferation, with cell division stopping abruptly thereafter (**Fig.1i**). A cluster of G3 cells followed G2 expansion and expressed secondary granule genes such as *Ltf*, *Camp*, and *Ngp* (**Fig.1e).** Neutrophil differentiation in the BM concluded with a more mature G4 population highly expressing *Mmp8*, a key granule protein for neutrophil-mediated host defenses, and Ccl6 which is important for neutrophil mobilization (**Fig.1d-e**). Finally, we measured the maturation score of each differentiating neutrophil population based on the expression of genes related to neutrophil differentiation and maturation (**Table S4**). G5 cells, which were mainly in the PB and peripheral tissues, were the most mature neutrophils, while G4 cells showed the highest maturation score among BM neutrophils (**Fig.1j**).

### A “sorting mechanism” for generating heterogeneous neutrophil granules

A well-accepted mechanism explaining neutrophil granule heterogeneity is “targeting by timing of biosynthesis” (Borregaard and Cowland, 1997; Borregaard et al., 2007; Cowland and Borregaard, 2016a), i.e., granule proteins synthesized at the same time in developing neutrophils will end up in the same granule without granule type-specific sorting. We examined the expression of various granule genes in differentiating neutrophils. Lactoferrin-positive granules are often defined as specific (secondary) granules, while lactoferrin-negative but gelatinase-positive granules are known as gelatinase (tertiary) granules. As expected, a major antibacterial specific granule protein, CAP-18 (*Camp* product) (Sorensen et al., 1997), was expressed in G2 and G3 cells when specific granules were formed. Surprisingly, this protein, which is not in the tertiary granules and secretary vesicles, was also expressed in G4 neutrophils containing tertiary granules and secretary vesicles.

Similarly, another specific granule protein, NGAL (*Lcn2* product), which co-localizes completely with lactoferrin (Kjeldsen et al., 1994), was also highly expressed in G4 neutrophils (**Fig.2a**). Thus, “targeting by timing of biosynthesis” may not explain all granule heterogeneity, and some granule proteins must be tagged to direct them to particular granules.

**Figure 2.**
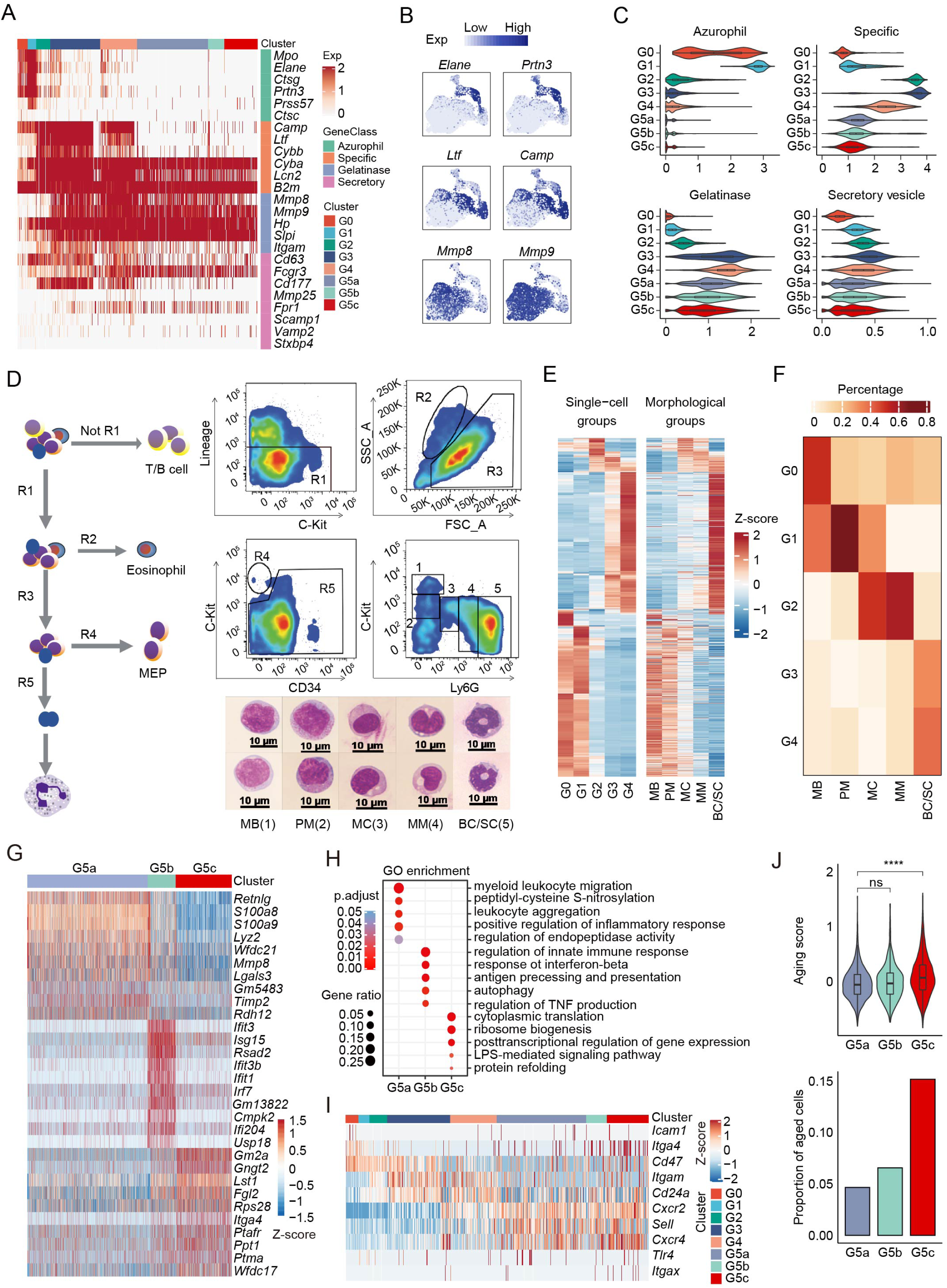
**(A-F) Transcriptional landscape of neutrophils along differentiation and maturation trajectories.** (A) Heatmap showing expression of neutrophil granule-related genes for all neutrophils. (B) Expression of six typical neutrophil granule genes. (C) Violin plots of azurophil score, specific score, gelatinase score, and secretory score for each cluster. (**D-F) scRNA-seq-defined differentiating neutrophil populations correlated with classical morphology-defined neutrophil subpopulations.** (D) FACS sorting and staining of five mouse BM neutrophil populations for bulk sequencing. Left: Gating diagram. R1 (CD4^-^ CD8a^-^CD45R/B220^-^Ter119^-^) was selected, R2 (eosinophil) and R4 (MEP) were excluded, and remaining R5 (neutrophils) were selected. Top right: Same R1-R5 from (left) but showing FACS gating of five detailed neutrophil subpopulations and the morphology of the sorted cells, among which: 1 was c-Kit^hi^Ly6g^neg^ (MB); 2 was c-Kit^int^Ly6g^neg^ (PM); 3 was c-Kit^neg^Ly6g^low^ (MC); 4 was c-Kit^neg^Ly6g^int^ (MM), and 5 was c-Kit^-^Ly6g^hi^ (BC/SC). Bottom right: Representative Wright-Giemsa staining of these populations (scale bar represents 10 μm); Data are representative of three independent experiments. (E) Heatmaps showing row-scaled expression of scRNA-seq-defined DEGs across averaged single-cell groups (left) and morphological groups (right). Only genes detected in both scRNA-seq data and bulk RNA-seq data are visualized. (F) Coefficient matrix showing deconvolution results of morphological bulk profiles. The 20 highest DEGs per single-cell group were selected as signatures for deconvolution. Each column is normalized by column sums. **(G-H) Transcriptional landscape of mature neutrophils in the peripheral blood and spleen.** (G) Heatmap showing row-scaled expression of the ten highest DEGs per cluster for G5a, G5b, and G5c neutrophils. (H) Gene ontology (GO) analysis of DEGs for each of the three G5 clusters. Selected GO terms with Benjamini-Hochberg-corrected P-values < 0.05 (one-sided Fisher’s exact test) are shown. **(I-J) Expression of neutrophil aging signatures.** (I) Heatmap showing row-scaled expression of aging-related genes for all neutrophils. (J) Top: Violin plot of aging score defined as weighted average Z-scores of aging-related genes for the three G5 clusters. Bottom: Proportions of aged cells in each G5 clusters.

Collagenase (*Mmp8* product), a key innate immunity and inflammation enzyme, was thought to be localized exclusively in specific granules (Murphy et al., 1977) and co-expressed with lactoferrin. However, at single-cell level, collagenase was always co-expressed with gelatinase, with the highest expression detected in G4 neutrophils and almost no expression in G2 cells highly expressing lactoferrin (**Fig.2a**), suggesting that collagenase should be considered a gelatinase rather than a specific granule protein.

### scRNA-seq-morphology correlates

We next compared scRNA-seq-defined neutrophil populations with the classic morphology-defined neutrophils. MBs, PMs, MCs, MMs, and mature neutrophils (including BCs and SCs) were isolated by FACS based on c-Kit and Ly6g expression (Satake et al., 2012), their identities confirmed morphologically (**Fig.2d**), and bulk RNA-seq performed. Most of the molecular signatures identified by scRNA-seq were also detected in bulk RNA-seq data (**Fig.2e and Fig.S2a**). To further dissect morphological heterogeneity, a regression-based deconvolution approach was applied on bulk expression profiles. MB were a mixture of G0/G1, PM were G1, MC were G1/G2, MM were G2, and BC/SC were G3/G4 (**Fig.2f**). MBs highly expressed the stem cell marker *Cd34* and translation-related genes such as *Eef1a1*, while the highest expression of primary granule-related genes was detected in the PM population **(Fig.S2a-b)**. MCs and MMs were highly proliferative neutrophils expressing high levels of cell cycle-related genes, which were significantly downregulated in mature neutrophils (**Fig.S2c**). BC/SC neutrophils expressed both G3 and G4 signature genes, confirming that this population is a mixture of G3 and G4 cells (**Fig.S2a)**.

### The PB and spleen contain three distinct neutrophil subpopulations

Three major neutrophil subpopulations were identified in the PB and spleen with 172 DEGs (**Table S3, Fig.2g**). Overall, BM and PB neutrophils were very different, implicating the microenvironment in driving neutrophil transcription. Similar to G4 BM cells, G5a cells highly expressed *Mmp8* and *S100a8* (**Fig.2g**) and genes related to neutrophil migration and inflammatory responses (**Fig.2h**). Interestingly, a group of G5b neutrophils expressed a set of interferon-stimulated genes (ISGs) such as *Ifit3* and *Isg15* (**Fig.S2d**). Although G5b appears to be a separate mature neutrophil subpopulation, they are not interferon-stimulated: unchallenged mice do not produce interferons, and unlike true interferon-induced ISG expression, many ISGs were not upregulated in the G5b population (e.g., *Irf3*, *lfitm6*) (**Fig.S2d**).

Trajectory analysis showed that G5a and G5b neutrophils gradually developed into G5c neutrophils (**Fig.S2e)** the latter showing the highest aging score (**Fig.2i-j**). Aging is a main mechanism that accounts for neutrophil heterogeneity (Adrover et al., 2016): aged neutrophils are smaller with fewer granules and granular multi-lobed nuclei, produce more NETs, and migrate out of capillaries less efficiently (Casanova-Acebes et al., 2013; Doerschuk et al., 1994; Kolaczkowska et al., 2015; Tanji-Matsuba et al., 1998). By applying a two-component Gaussian mixture model, we further identified 15% of G5c neutrophils as aged, significantly higher than in G5b or G5a populations (**Fig.2j**). Although G5c cells appeared to more aged, the mitochondrial UMI percentage was not elevated in G5c cells, indicating continued viability in the PB and spleen (**Fig.S2f)**. G5c marker genes were significantly enriched for ribosome biogenesis, cytoplasmic translation, post-transcriptional regulation, and LPS-mediated signaling pathways by GO analysis (**Fig.2h**). These results agree with recent studies showing that aged neutrophils are highly functional and may exhibit a higher phagocytic activity than non-aged neutrophils (Kolaczkowska, 2016; Uhl et al., 2016).

### The pathogen clearance machinery is continuously and gradually built during neutrophil differentiation, maturation, and aging

We next assessed the expression of genes critical to several typical neutrophil-mediated microbial clearance functions. Phagocytosis, chemotaxis, and neutrophil activation scores increased drastically during the early stages of granulopoiesis, peaked at G3, and remained relatively stable thereafter (**Fig.S3a-c**). In neutrophils, phagocytic NADPH oxidase is responsible for pathogen-induced ROS production and ROS-mediated pathogen killing. Similarly, “NADPH oxidase score”, calculated based on the expression of the seven NADPH oxidase-related genes, increased during G0 to G1 and G2 transition, peaked at G3, and then decreased by 20% in mature neutrophils (**Fig.S3d**). However, the dynamics of the oxidase complex subunits varied through neutrophil differentiation (**Fig.S3e**): *Cybb* (gp91-phox) was maximally expressed in G2 and G3 cells and abruptly downregulated thereafter; and *Ncf2* was upregulated in G3, peaked in G4 cells, and then remained stable. Sequential subunit expression ensures maximal stimulation-triggered NADPH oxidase activation at the later stages of neutrophil maturation and minimal activation in immature neutrophils in the BM. Collectively, this tightly orchestrated neutrophil maturation program ensures that maximum neutrophil activation only occurs when neutrophils are fully mature and mobilized to the PB so that innate host defense execution is effective and safe. Notably, genes related to mitochondria-mediated ROS production were significantly downregulated during neutrophil maturation, further supporting that neutrophil ROS production is mainly mediated by phagocytic NADPH oxidase (**Fig.S3f**).

Mature neutrophils derive energy mainly from glycolysis (Borregaard and Herlin, 1982). However, metabolism-related genes **(Fig.S3g)**, including those related to glycolysis **(Fig.S3h)** were downregulated in mature neutrophils. Similarly, genes related to glucose transportation were also not upregulated in mature neutrophils **(Fig.S3i)**. These data suggest that glycolysis-dominant metabolism in neutrophils is likely to be driven by post-transcriptional or/and post-translational mechanisms rather than transcriptional upregulation of related genes.

### Organ-specific transcriptome features

Compared to early-stage maturing BM neutrophils, most neutrophils in the PB and SP were a mature G5 population with similar gene expression patterns (**Fig.S3j**). PB neutrophils expressed chemokines (e.g., *Il1b*), ribosomal genes (e.g., *Rps27rt*), and genes related to neutrophil activation and function (e.g., *Csf3r, Fgl2*). Notably, SP neutrophils highly expressed the transcription factors *Fos* and *Junb*, chemokine (C-X-C motif) ligand 2 (*Cxcl2/Mip2a*), HLA class II histocompatibility antigen gamma chain (*Cd74*), and certain myeloid transcription factors (e.g., *Cebpb*) (**Fig.S3j**). KEGG enrichment analysis revealed PB neutrophils were enriched for genes related to malaria and African trypanosomiasis, while SP neutrophils were more enriched for genes related to leishmaniasis and *Yersinia* infection, suggesting that PB and SP neutrophils may play distinct roles in combating different infections (**Fig.S3k**). SP neutrophils also expressed T cell differentiation, IL-17 signaling, TNF signaling, and antigen processing and presentation-related genes, suggesting a role for splenic neutrophils in adaptive immunity. Indeed, a recent study indicated that neutrophils can adopt some APC function (Vono et al., 2017), and splenic B cell-helper neutrophils participate in immunoglobulin diversification and production in the marginal zone (Puga et al., 2011).

### Unexpected complexity in neutrophil mobilization

To further elucidate the relationship between the new neutrophil subpopulations, we traced cell fate and reconstructed cell lineage direction using the recently developed RNA ‘velocity’ approach (La Manno et al., 2018b) (**Fig.3a**). Consistent with Monocle (**Fig.1h**), BM maturation (from G2 to G4) followed a single main branch without significant division, with G3 bearing long vectors and indicating a strong tendency to progress to G4 (**Fig.3a**). Neutrophils are thought to mobilize from the BM to the PB only at full maturity. Interestingly, the trajectory of a significant number of G3 neutrophils was toward the peripheral G5a population, suggesting mobilization of G3 cells to the PB or tissue without first undergoing full G4 maturation (**Fig.3a**). BM G4 population split into: (i) the peripheral G5a population, and (ii) the ISG-related G5b population without entering the G5a stage. Thus, although G5a and G5b were most similar (**Fig.1g**), they are two separate and independent PB neutrophil populations, with G5a derived from BM G3 and G4 cells and G5b derived solely from G4 cells. G5a to G5b conversion was rarely detected in the PB (**Fig.3a**). Lastly, G5c cells were firmly at the end of neutrophil maturation and differentiation, showing the highest apoptosis scores (**Fig.3b)** and proportion of apoptotic cells (∼30%; **Fig.3c**). There was also significant apoptosis in G5a and G5b cells (**Fig.3c),** suggesting that death programs can be independent of maturation.

**Figure 3.**
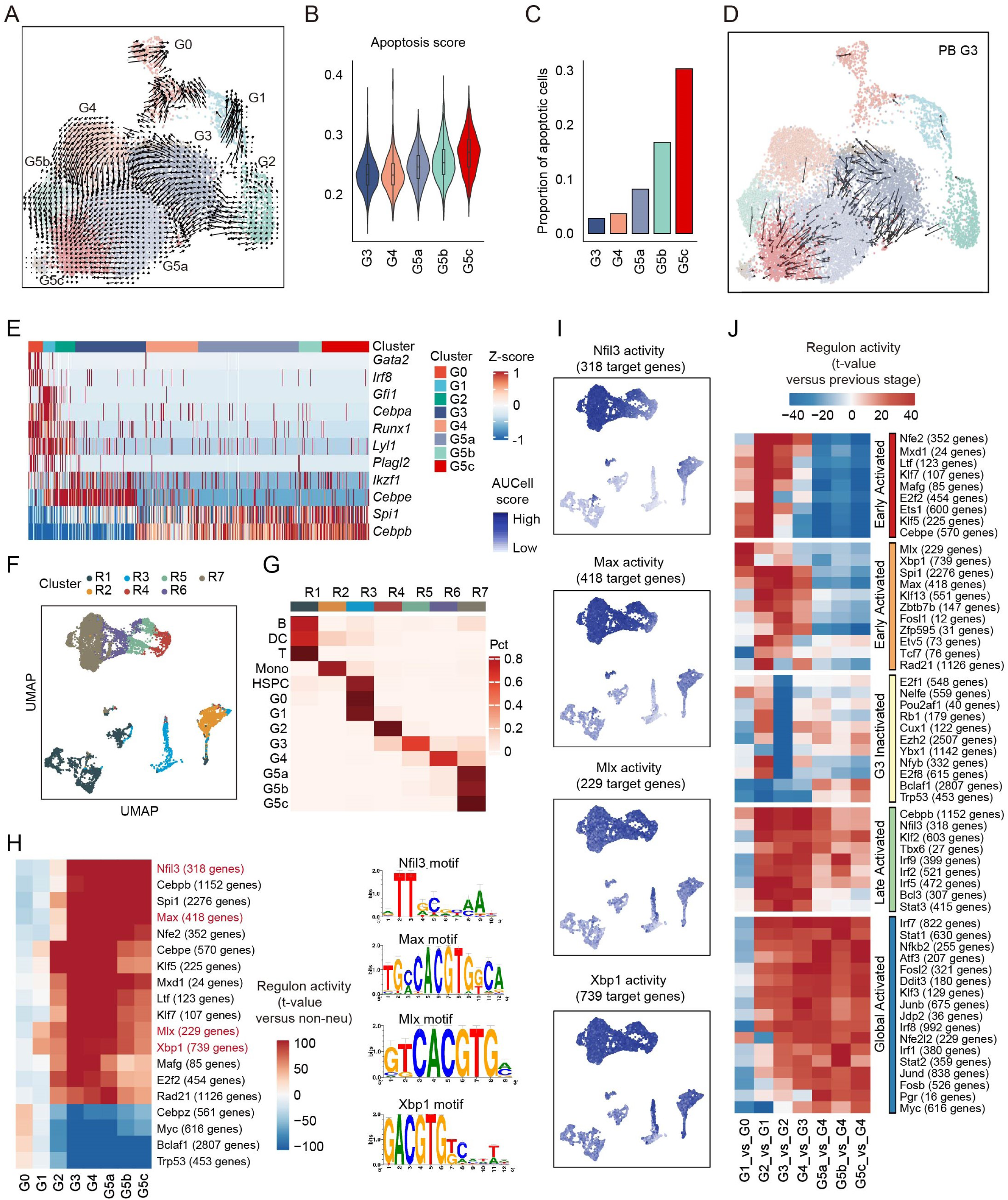
**(A-D) The origin and inter-relationship of neutrophil subpopulations.** (A) Velocity analysis reveals the origin and inter-relationship of neutrophil subpopulations. Velocity fields were projected onto the UMAP plot. (B) Violin plot of apoptosis scores (GO:0097193) for G3-G5 clusters. (C) Proportions of apoptotic cells in each cluster identified by a two-component Gaussian mixture model. (D) As in (A) but only of G3 neutrophils originating from PB. **(E-J) The formation of neutrophil subpopulations is driven by both known and a large set of uncharacterized transcription factors.** (E) Heatmap showing row-scaled gene expression of TFs known to be involved in granulopoiesis and neutrophil function. (F) UMAP of the regulon activity matrix of neutrophils and 7209 non-neutrophils under normal conditions. K-means clustering was performed on the first 20 principal components (PCs) of the regulon activity matrix with cluster number *k* = 7. Each cell is assigned the color of its K-means cluster. (G) Confusion matrix showing the percentage overlap of Seurat transcriptome-based clusters with K-means regulon-based clusters. (H) Heatmap of the t-values of regulon activity derived from a generalized linear model of the difference between cells from one neutrophil subpopulation and cells from other non-neutrophil populations. Only regulons with at least one absolute t-value >100 are visualized. Previously uncharacterized neutrophil-specific transcription factors are marked in red with binding motif shown on the right. (I) Activities of the four newly identified neutrophil-specific regulons. (J) As in (H), but t-values representing activity change between the current developmental stage and the previous one. Only regulons with at least one absolute t-value >40 are visualized. Regulons are hierarchically clustered based on activation pattern (red and orange: early-activated, yellow: G3-inactivated, green: late-activated, blue: global-activated).

A small number of immature neutrophils also circulate, which are thought to be derived from accidental release of cells closest to maturation (Broxmeyer, 2008; Furze and Rankin, 2008; Suratt et al., 2001). Both the G3 and G4 populations are differentiating neutrophils that mainly exist in the BM. However, we only detected G3 cells in the periphery of healthy mice (5% of PB and 6% of SP neutrophils; **Fig.1c**). During BM granulopoiesis, only G3 neutrophils seem to be able to escape from the BM niche, migrate into PB, and travel to other organs. PB and BM G3 cells consistently overlapped on velocity analysis, with some falling into the PB G5a cluster **(Fig.3d**). Further, PB G3 cells directly differentiated into G5a without going through G4, consistent with the absence of G4 cells in the PB and SP (**Fig.3d**).

### Both known and many uncharacterized transcription factors drive neutrophil subpopulations

We next sought to characterize transcription factor (TF) dynamics across neutrophil differentiation and maturation, since tightly regulated transcriptional programs are likely to dictate neutrophil populations (Monticelli and Natoli, 2017; Theilgaard-Monch et al., 2005). We first examined the expression of TFs known to be involved in granulopoiesis and neutrophil function (**Fig.3e**). Genes related to stem cell maintenance and early lineage commitment such as *Gata2*, *Irf8*, and *Runx1* were highly expressed in the G0 population. Genes highly expressed in G1 included *Gfi1* and *Cepba*, which play essential roles in neutrophil development (Avellino and Delwel, 2017; Guo et al., 2012; Hock et al., 2003; Zhang et al., 1997), strongly suggesting that specific neutrophil lineage commitment occurs during G1. Cebpa has been implicated in cessation of cell proliferation during granulopoiesis (Porse et al., 2001; Slomiany et al., 2000). However, *Cebpa* expression was highest in proliferating G0 and G1 cells and then drastically declined in more mature cells, with the lowest expression found in the non-dividing G3 population, making it unlikely that Cebpa directly ceases proliferation of maturating neutrophils. Similarly, Cebpe is another terminal neutrophil differentiation factor (Bjerregaard et al., 2003; Gombart et al., 2003; Gombart et al., 2001; Khanna-Gupta et al., 2007; Larsen et al., 2014; Paul et al., 2016; Verbeek et al., 2001; Yamanaka et al., 1997) implicated in promoting cell cycle arrest (Gery et al., 2004; Nakajima et al., 2006), but *Cebpe* transcription was highest in proliferating G2 cells (**Fig.3e**). Thus, the transcriptional switch ending cell proliferation stage and initiating terminal differentiation is likely to be controlled by TFs and networks other than *Cebpa* and *Cebpe*.

To assess specific global gene regulatory networks associated with neutrophil maturity, we applied SCENIC analysis (Aibar et al., 2017). Based on regulon activity, cells were projected onto a low-dimensional subspace, reproducing similar developmental trajectories to those of the Seurat projections (**Fig.3f**). There was high consistency between Seurat clusters and SCENIC clusters (**Fig.3g**). HSPCs, G0, and G1 formed an aggregate cluster, suggesting a shared “primitive” regulatory state, whereas the other four non-neutrophil populations had dramatically different TF networks. To further dissect the regulatory differences between neutrophils and other cell types, we compared regulon activities from each neutrophil group versus all non-neutrophil populations using a generalized linear model (Lambrechts et al., 2018). This identified 19 neutrophil-specific networks, including previously reported TFs such as *Cebpe, Spi1*, and *Klf5* (**Fig.3h and Table S5**). Importantly, this analysis also identified four new regulons, *Nfil3*, *Max*, *Mlx*, and *Xbp1*, which are closely related to the expression of neutrophil-specific genes (**Fig.3i**).

We next examined the regulatory events responsible for transitioning between consecutive neutrophil differentiation stages (**Fig.3j**). Coarse-grained clustering revealed at least five regulon groups with distinct activation patterns, including two early-activated, one late-activated, one globally-activated, and one specifically-inactivated after G2. While many TF networks like *Cebpe, Ets1, Klf5, Rad21,* and *E2f2* contributed to neutrophil commitment, changes in *Xbp1* and *Mlx* networks were specifically associated with G0/G1 transition. Additionally, the dramatic loss of regulatory networks such as *E2f1*, *Nelfe*, and *Rb1* indicated a potential functional change between G2 and G3. As well as TFs known to be essential for neutrophil differentiation and maturation, we also identified numerous novel regulons responsible for transitioning, providing a foundation for further studying the molecular basis of neutrophil maturation and heterogeneity.

### Bacterial infection primes neutrophils for augmented functionality without affecting overall heterogeneity

We next investigated how bacterial infection affected neutrophil subpopulations (**Fig.S4a**), including in the liver and peritoneal cavity (**Fig.S4a-c)**. At the same sequencing depth, the gene number and total UMIs both increased in neutrophils isolated from the PB, SP, and PB of *E. coli*-challenged mice compared to control mice, indicating elevated transcriptional activity during bacterial infection (**Fig.S4d-e**). Leveraging the fact that cells in each subpopulation from unchallenged mice share common signature genes, we were able to identify every population in challenged mice using a well-accepted data integration method (Stuart et al., 2019) (**Fig.4a-b**) and validated independently through unsupervised dimension reduction at both transcriptome and transcription regulatory network levels (**Fig.4a-b** and **Fig.S6a**). The expression of signature molecules (**Fig.4c**), NADPH oxidase components (**Fig.S4f**), and granular proteins (**Fig.S4g**) remained remarkably consistent after *E. coli* challenge. Thus, the identity of each neutrophil population was maintained during acute bacterial infection and the signature genes still successfully determined neutrophil identity under inflammatory conditions (**Fig.4c**).

**Figure 4.**
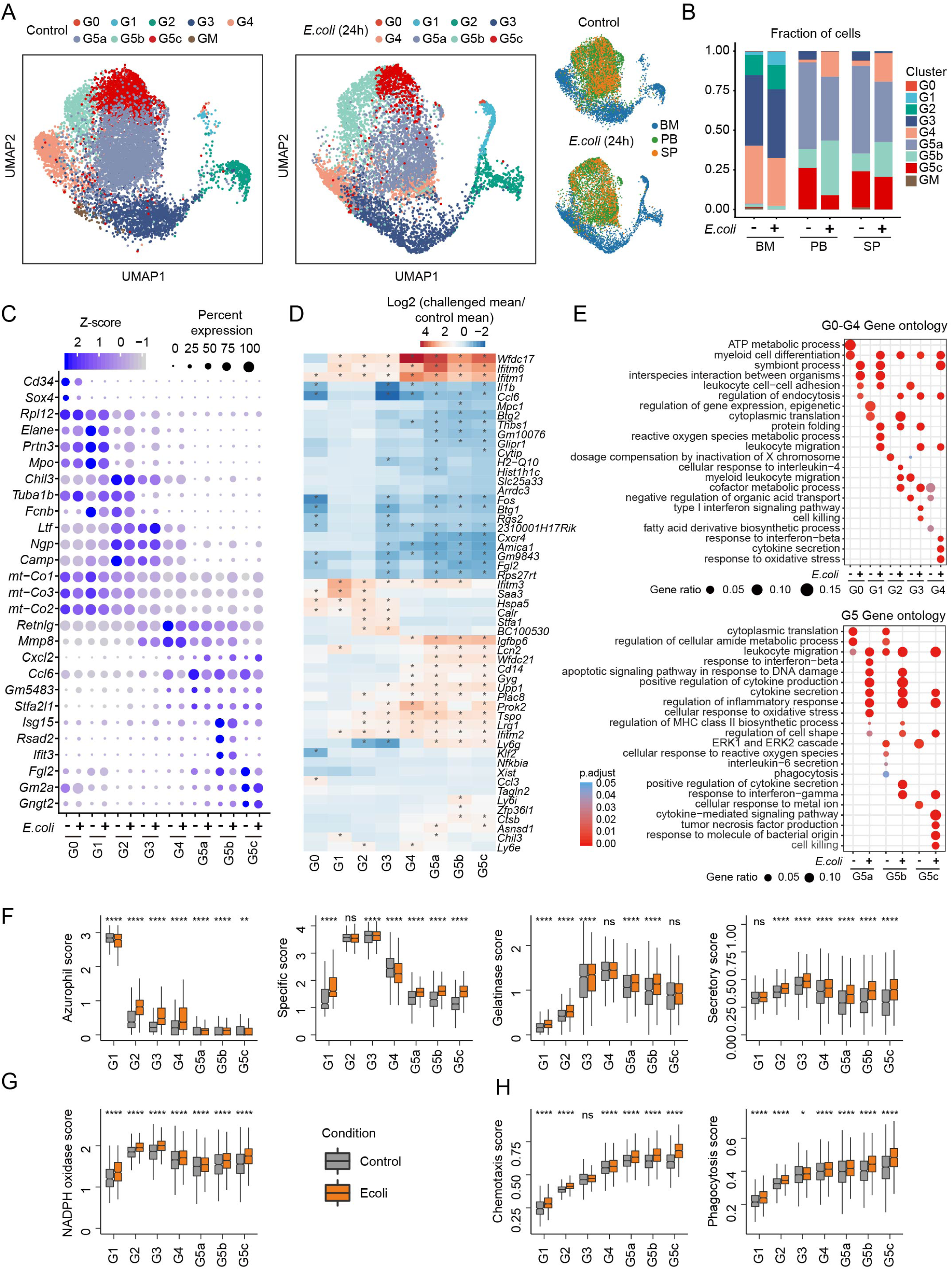
Bacterial infection primes neutrophils for augmented functionality without affecting their overall heterogeneity. (A) Comparison of control and *E. coli*-challenged neutrophils originating from BM, PB, and SP. All neutrophils under both control and *E. coli-*challenged conditions (13,687 cells) are projected together by UMAP but are displayed separately by experimental condition. (B) Comparisons of neutrophil composition between control and *E. coli* challenge in BM, PB, and SP before and after *E. coli* challenge. (C) Dot plot showing scaled expression of signature genes for each cluster before and after *E. coli* challenge colored by average expression of each gene in each cluster scaled across all clusters. Dot size represents the percentage of cells in each cluster with more than one read of the corresponding gene. (D) Heatmap showing log2(fold-change) in gene expression of the representative cluster-based DEGs between control and *E. coli*-challenged neutrophils. The asterisks mean log2(fold-change)>1 in corresponding cells. (E) GO analysis of cluster-based DEGs between control and *E. coli*-challenged neutrophils. Selected GO terms with Benjamini-Hochberg-corrected P-values < 0.05 (one-sided Fisher’s exact test) are shown. (F-H) Comparisons of functional scores between control and *E. coli*-challenged neutrophils for each cluster. (F): Granule scores; (G): NADPH oxidase complex score; (H): Chemotaxis score and phagocytosis score. ns, P>0.05; *, P <=0.05; **, P <= 0.01; ***, P < 0.001, ****, P <= 0.0001. student’s t-test.

Despite the overall subpopulation stability during bacterial infection, infection up- and downregulated numerous genes in each neutrophil subpopulation (**Fig.4d**). Several cytokines involved in early leukocyte recruitment during infection such as *Il1b* and *Ccl6* were downregulated in neutrophils from *E. coli*-challenged hosts. In differential gene expression analysis (**Fig.S5a and Table S6**), DEGs in G0 and G1 cells were also preferentially involved in regulating immune effector processes and ROS metabolism, respectively, suggesting that immune adaptation to bacterial infection could occur as early as within early progenitor cells (**Fig.4e and Fig.S5b**). Under steady-state conditions, protein translation genes were highly expressed in G0 and G1 cells (**Fig.1f**), consistent with their highly proliferative nature. During bacterial infection, these genes and genes related to protein folding were further upregulated in G2 and G3 cells, presumably to meet the elevated protein needs of functionalized neutrophils (**Fig.S5b)**. In relatively mature G4 and G5 neutrophils, bacterial infection triggered significant upregulation of cytokine production and secretion genes (**Fig.4e and Fig.S5b**).

Interestingly, genes related to response to interferon-beta were preferentially upregulated in G5a cells, while genes related to T cell proliferation and activation were upregulated in G5b cells (**Fig.S5b)**, suggesting that these two neutrophil populations play different roles in bacterial infection. Finally, in bacteria-challenged hosts, neutrophil functions related to bactericidal activities including synthesis of granular proteins (**Fig.4f**), NADPH oxidase complex (**Fig.4g**), phagocytosis, and chemotaxis (**Fig.4h**) were all upregulated. Thus, during bacterial infection, core neutrophil subpopulations are maintained but genes related to pathogen clearance are upregulated at each stage of neutrophil maturation to maximize host defenses.

We also examined whether bacterial infection had a universal impact on transcriptional regulatory networks across neutrophil populations. Overall, there was a coherent drift in gene regulatory network activities in each subpopulation after bacterial challenge (**Fig.S6a**), perhaps driven by upregulation of defense-response-associated TF networks like *Irf7* and downregulation of metabolic TF networks like *Foxp1* and *Ctcf* (**Fig.S6b**). Interestingly, we also identified TF networks (e.g. *Fos* and *Atf4*) showing different responses in immature (upregulated) and mature (downregulated) populations (**Fig.S6b**). These networks were gradually activated from G1 to G5 under normal conditions (**Fig.S6c and Table S5)**, and their target genes were enriched for a variety of processes including signaling, biosynthesis, and transcriptional regulation. Presumably, bacterial challenge accelerates neutrophil maturation by upregulating these networks at earlier stages and thus re-allocates cellular resources to defense responses by downregulating these networks at later stages.

### The liver displays a distinct extramedullary granulopoiesis program during bacterial infection

The liver is a major contributor to embryonic and fetal hematopoiesis but only a minor contributor to adult hematopoiesis (Mafra et al., 2019). In healthy mice, neutrophil numbers in the liver were so low that we could not collect enough Gr1^+^ neutrophils for scRNA-seq. Extramedullary granulopoiesis during infection generates sufficient mature neutrophils for pathogen clearance (Kim, 2010) and enable scRNA-seq. Both the percentage and total number of neutrophils increased significantly in the liver after *E. coli* challenge (**Fig.S6d)**. In terms of liver neutrophil composition (**Fig.S6e**), the granulopoiesis trajectory in the liver was more obvious than that in the spleen, where there was almost completely overlap with PB neutrophils (**Fig.4a**) and the percentage of G0, G1, and G2 cells were significantly lower (**Fig.S6f**). Interestingly, only about half of liver neutrophils overlapped with PB or BM neutrophils (**Fig.S6e**), the other half (mainly G3 and G4 neutrophils) forming a distinct liver-specific population (**Fig.S6e**). The origin of the liver G0-G2 cells remains elusive. HSPCs are mobilized from the BM to the peripheral tissue where they may contribute to extramedullary granulopoiesis. Alternatively, they can be generated locally from more primitive cells. A previous study suggested that extramedullary hematopoiesis in the adult mouse liver is associated with specific hepatic sinusoidal endothelial cells (Cardier and Barbera-Guillem, 1997).

### The ISG-related G5b neutrophil population exists in both humans and mice and expands during infection

The percentage of G5b cells increased significantly in *E. coli*-challenged hosts (**Fig.4b**). We investigated the distribution of G5b neutrophils in the spleen using a laser scanning cytometer (LSC), co-staining spleen tissue sections with an anti-S100a8 antibody and an anti-Ifit1 antibody to identify G5b neutrophils (**Fig.5a**). Under normal conditions, S100a8^+^ neutrophils were uniformly distributed in the red pulp, while G5b (S100a8^+^ Ifit1^+^) cells were preferentially subcapsular. After *E. coli* challenge, overall number of neutrophils in the spleen increased significantly, as did the percentage and number of G5b (**Fig.5b**), which were still preferentially subcapsular, their specialized location further demonstrating the uniqueness of this subpopulation. Although multiple ISGs (e.g. *Ifitm1*) were upregulated in basically all neutrophil subpopulations after bacterial stimulation (**Fig.5c**), many ISGs such as *Isg15* and *Oas2* which are specifically expressed in G5b were not upregulated, suggesting that ISG-related G5b expansion was not due to bacteria-induced ISG expression. We next examined whether this G5b population was also present in human blood by scRNA-seq of sorted human PB neutrophils from a healthy donor. Unsupervised clustering revealed three major neutrophil populations, one of which was a human G5b (hG5b) population accounting for 35% of the PB neutrophils (**Fig.5d-e and Table S7**). Similar to mouse G5b neutrophils, hG5b only expressed a subset of ISGs (**Fig.5f**).

**Figure 5.**
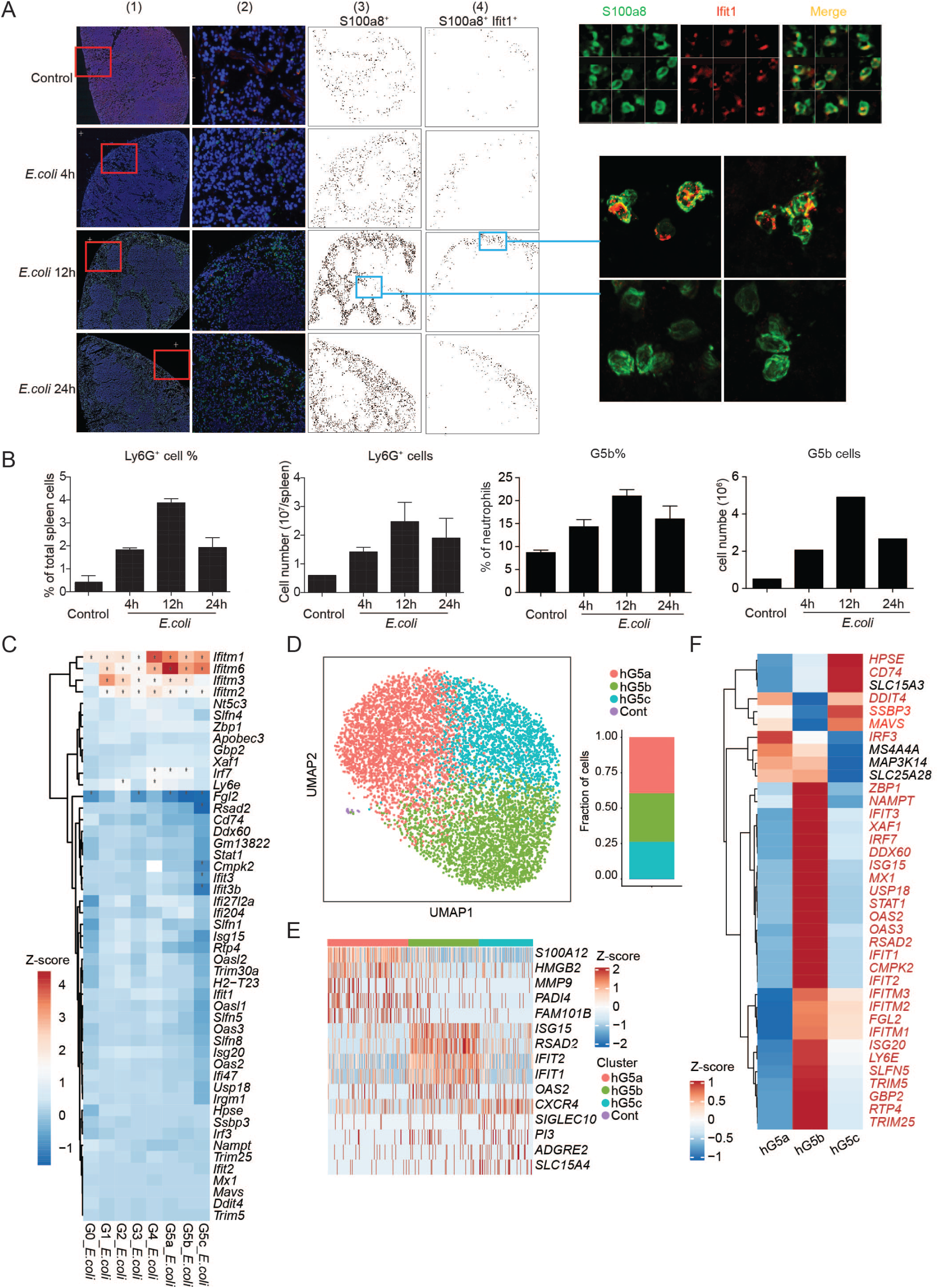
**The ISG-expressing neutrophil population is present in both humans and mice and expands during bacterial infection.** (A) Quantitative image analysis of the spatial distribution of G5b in whole spleen sections. Left: Laser scanning cytometry (LSC) image analysis of the whole spleen. (1) Low-magnification image of axial spleen cryosection immunostained for DAPI (blue), S100a8 (green), and Ifit1 (red). (2) Representative images of spleens from the selected region from (1). (3) Localization diagram of S100a8^+^ cells in the whole spleen from (1). Cells gated positive based on the fluorescence intensity in the S100a8 channel. Each dot represents a single cell. (4) Localization diagram of S100a8^+^Ifit1^+^ (G5b) cells gated from (3). Each dot represents a single cell. Right: LSC images (top) and confocal images (bottom) of representative G5b neutrophils. (B) Quantification and relative frequency of spleen Ly6g^+^ cells (left, measured by FACS) and spleen G5b cells (right, S100a8^+^ Ifit1^+^, measured by LSC) at different time points after *E. coli* infection. Results are the mean ± SD of three independent experiments. (C) Heatmap showing log2 fold change in expression of 49 ISGs before and after *E. coli* challenge for each cluster. Genes are marked with an asterisk if their expression changed significantly as identified by a Student’s t-test (Bonferroni-corrected P-value < 0.05). (D) UMAP of neutrophils from human peripheral blood (PB) colored by cluster identity. The fraction of cells in each cluster is displayed on the right. (E) Heatmap showing row-scaled expression of the five highest DEGs (Bonferroni-corrected P-values < 0.05, Student’s t-test) per cluster for all hG5 neutrophils. (F) Heatmap showing expression of 37 ISGs for the three human neutrophil clusters. Genes marked in red are conserved across mouse and human.

### Bacterial infection accelerates G1 cell division and post-mitotic maturation without altering overall neutrophil differentiation

Neutrophil populations significantly expand during bacterial infection (Kwak et al., 2015; Manz and Boettcher, 2014). However, the neutrophil differentiation and maturation trajectory was largely maintained in *E. coli*-challenged mice (**Fig.6a**). The overall stability of the neutrophil differentiation program after bacterial infection was also demonstrated by correlation of SCENIC transcription regulatory networks in control and challenged samples (**Fig.6b**). We next explored the mechanisms elevating neutrophil production during acute bacterial infection. While both G1 and G2 cells are proliferative, the proliferation score increased only in the G1 population (**Fig.6c**), as were genes related to G2/M phase progression (Tirosh et al., 2016a), while genes related to S phase progression were paradoxically reduced in G2 cells during acute infection (**Fig.6d**). The accelerated cell proliferation seen during emergency granulopoiesis is therefore likely to be mediated by enhanced cell proliferation in the G1 population. This was also supported by the more drastic expansion of G1 rather than G2 population during acute infection (**Fig 4b**).

**Figure 6.**
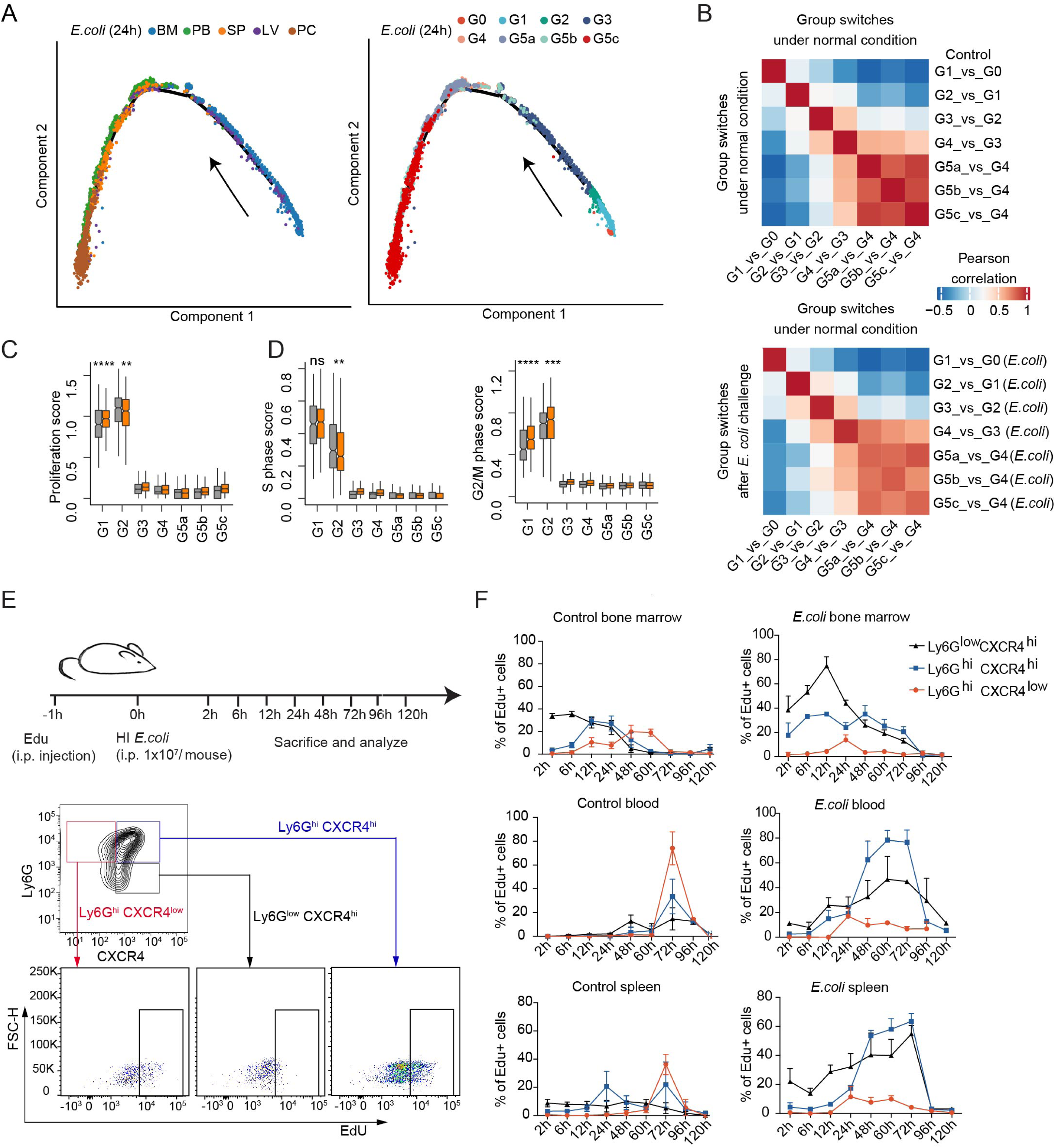
**Bacterial infection accelerates G1 cell division and post-mitotic maturation without altering overall neutrophil differentiation programs.** (A) Monocle trajectories of *E. coli-*challenged neutrophils colored by sample origin (left) and cluster identity (right). Each dot represents a single cell. Cell orders are inferred from the expression of the most variable genes across all cells. The trajectory direction was determined by biological prior. (B) Correlation matrices of t-values for regulon activity change during each group transition event under normal conditions (top) or after *E. coli* challenge (bottom). For each group transition event after challenge, the direct comparison to all normal transition events is demonstrated (bottom). (C-D) Comparisons of proliferation score (C), S-phase score, and G2M score (D) between control and *E. coli*-challenged neutrophils for each of the 8 clusters. (E-F) *In vivo* EdU incorporation assay. (E) Top: Schematic. Bottom: Gating strategy of the three neutrophil subpopulations: immature (Ly6g^low^Cxcr4^hi^, black), intermediate (Ly6g^hi^Cxcr4^hi^, blue), and mature (Ly6g^hi^Cxcr4^low^, red) neutrophils. (F) *In vivo* EdU proliferation assay of neutrophil subsets in BM, PB, and SP at sequential time points with or without *E. coli* challenge. Data are represented as percentages of EdU^+^ cells in the corresponding gated subpopulation. Results are the mean (±SD) of three independent experiments.

During bacterial infection, the G3 and G4 pool must increase to produce more mature neutrophils. Accelerated progenitor cell proliferation also suggests increased input to the post-mitotic G3 and G4 cells. Nevertheless, the proportions of G3 and G4 cells were not increased in the BM of *E. coli*-challenged hosts (**Fig.4b**), indicating that post-mitotic maturation may be accelerated in the BM. To test this hypothesis, we labeled dividing cells with 5-ethynyl-2’-deoxyuridine (EdU) and tracked these cells post-mitotically in the BM and PB (**Fig.6e**). Based on the surface expression of Ly6G and Cxcr4 in differentiating neutrophils (Eash et al., 2010; Evrard et al., 2018; Giladi et al., 2018), we separated relatively immature (Ly6G^low^Cxcr4^hi^), intermediate mature (Ly6G^hi^Cxcr4^high^), and mature (Ly6G^hi^Cxcr4^low^) neutrophils, and examined the emergence of EdU-labeled cells in these subpopulations at different time points. In the BM of unchallenged mice, EdU-labeled cells entered the immature neutrophil stage after 2 h, intermediate maturation after 12 h, and became mature neutrophils at 48 h. The duration of each stage was about 24 h, and EdU^+^ cells appeared in the PB and SP after 72 h (**Fig.6f**). In *E. coli*-challenged hosts, the percentage of Ly6G^hi^Cxcr4^low^ cells reduced significantly, and Ly6G^low^Cxcr4^hi^ and Ly6G^hi^Cxcr4^hi^ cells predominated in both the BM and PB. These neutrophils mobilized to the periphery following a similar dynamic pattern but over only two rather than three days, a drastic reduction in the post-mitotic neutrophil maturation period in infected hosts (**Fig.6f**).

### Bacterial infection reprograms the neutrophil population structure and dynamic transitions between subpopulations

Bacterial infection did not alter the overall identity of each neutrophil subpopulation but instead resulted in dynamic changes in each neutrophil population. In the BM, the proportion of G1 cells increased, indicating elevated proliferation of myeloid progenitors **(Fig.4b and Fig.7a)**. The percentage of BM G2 cells remained the same, suggesting balanced influx from G1 cells and transformation from G2 to G3 cells. In velocity analysis **(Fig.7b and Fig.S7a-b)**, the obvious transformation from G3 to G5a cells under homeostatic conditions was suppressed during infection, and G3 cells in infected hosts predominantly differentiated to G4 cells **(Fig.7b)**. G4 cells decreased from 38% to 30% in the BM but significantly increased in the PB and SP (**Fig.4b and Fig.7a**). PB G4 cells were mainly derived from the BM G3 cells (**Fig.7b)**. Additionally, infection significantly suppressed the G5a and G5b to G5c transition, leading to a smaller G5c population in *E. coli* challenged hosts (**Fig.7b)**. To assess neutrophil heterogeneity at the site of infection, neutrophils were extracted from the peritoneal cavity (PC) (**Fig.7c-d)**. Very few immature cells were detected, with >97% of neutrophils being mature (G5a-c). Interestingly, although G5c cells accounted for only about 25% of PB neutrophils before bacterial challenge and <10% PB neutrophils after bacterial challenge, >45% of PC neutrophils in challenged mice were G5c cells (**Fig.7d)**, suggesting that these cells may possess higher trans-endothelial migration capability than G5a or G5b cells. Supporting this hypothesis, the number of circulating G5c cells drastically reduced from 26% to 8.2% after bacterial challenge **(Fig.4b).**

**Figure 7.**
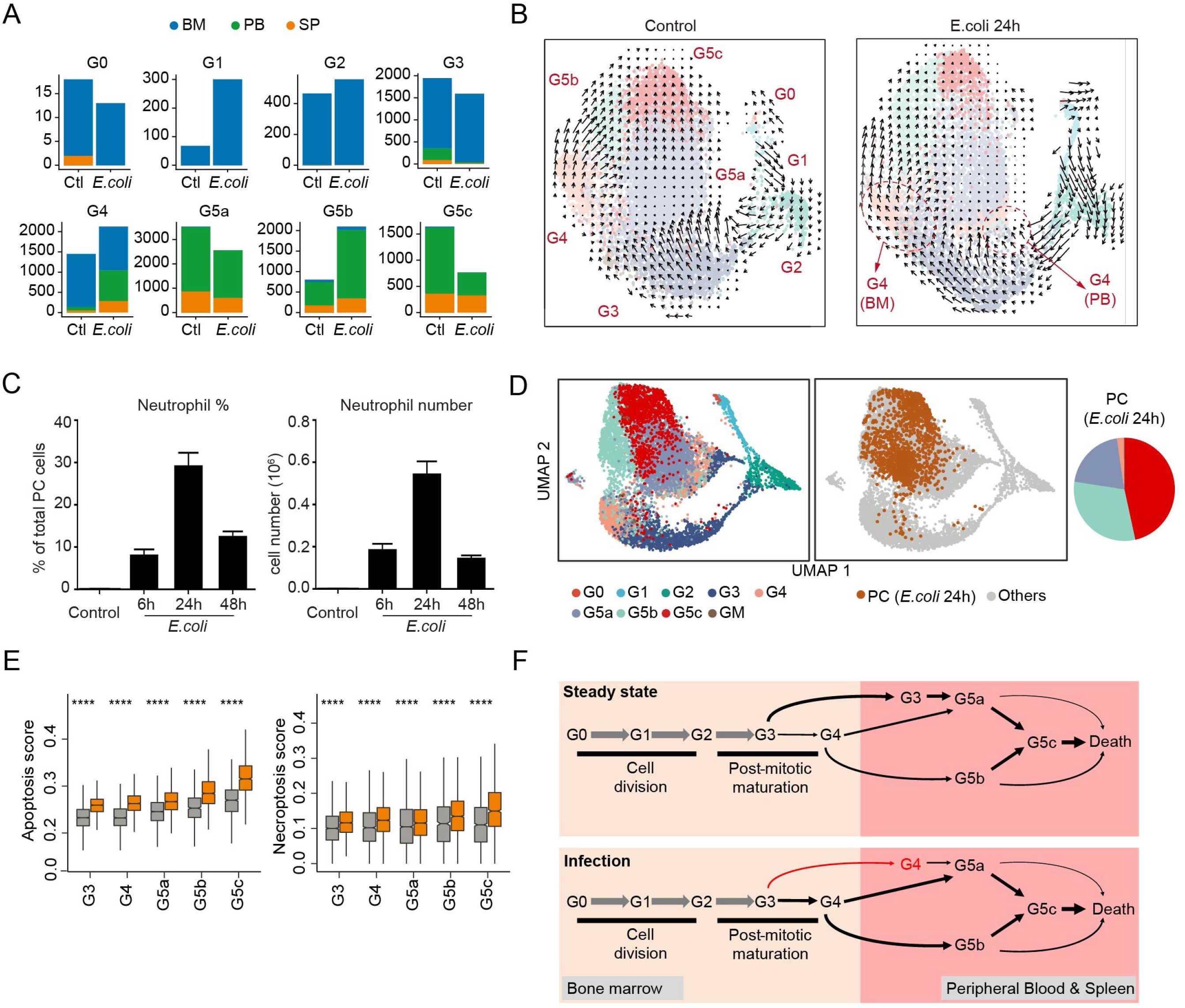
**Bacterial infection reprograms the structure of the neutrophil population and the dynamic transition between each subpopulation.** (A) Comparisons of organ distributions of each neutrophil subpopulation before and after *E. coli* challenge, measured by cell number. (B) Comparison of neutrophil dynamics (velocity field projected on the UMAP plot) before and after *E. coli* challenge. (C) Neutrophil proportion and cell number in the peritoneal cavity measured at different time points after *E. coli* challenge. Results are the mean (±SD) of three independent experiments. (D) Left: UMAP of *E. coli*-challenged neutrophils from BM, PB, SP, liver (LV), and peritoneal cavity (PC) colored by cluster identity. Right: PC cells are highlighted in the UMAP plot. Proportions of each neutrophil clusters in LV were shown. (E) Comparisons of apoptosis score and necroptosis score between control and *E. coli*-challenged neutrophils for G3-G5 clusters. ****, P ≤ 0.0001. Student’s t-test. (F) Dynamic transition between each subpopulation under steady-state and bacterial infection conditions. We cataloged differentiating and mature mouse neutrophils in an unbiased manner using single-cell RNA sequencing. Committed neutrophils include two proliferating subsets, two post-mitotic maturing subsets, and three functional mature subsets. Under homeostatic condition, the G5a and G5b cells in the PB arise from BM G4 and G3 cells, respectively. The transformation from G3 to G5a cells was suppressed during infection and G3 cells in infected hosts predominantly differentiated to G4 cells.

Infection delays neutrophil death (Luo and Loison, 2008). Paradoxically, genes related to apoptosis or necroptosis were significantly upregulated in every neutrophil subpopulation in *E. coli* challenged mice (**Fig.7e)**, indicating that the delayed neutrophil death in infected hosts is mainly determined by the activation of apoptotic factors and pathways rather the level of the related proteins.

## Discussion

A complete mechanistic understanding of neutrophil differentiation and function requires detailed knowledge of the gene expression alterations driving progression from one cellular state to the next or accompanying a particular functional population. Neutrophils are transcriptionally active (Silvestre-Roig et al., 2016; Zhang et al., 2004), and many aspects of transcriptional dynamics, including epigenetic regulation, are well documented (Grassi et al., 2018; Rönnerblad et al., 2014; Schuyler et al., 2016). Nevertheless, expression profiling of bulk, heterogeneous cell populations captures average expression states that may not represent any single cell. Here we used single-cell RNA sequencing to investigate the transcriptional landscape of neutrophil maturation and fate decisions under steady-state conditions and during bacterial infection at the single-cell level. Unbiased profiling of immature and mature neutrophils in the bone marrow, peripheral blood, and spleen provides new opportunities for exploring the transcriptional landscape of neutrophil development, function, and organ distribution.

It is well accepted that neutrophils are an inhomogeneous population. However, the presence of defined neutrophil subpopulations and whether they serve specific functions are a matter of debate. A common argument is that neutrophil heterogeneity may simply represent different neutrophil developmental stages, activation states, and microenvironments. Additionally, the specific function of circulating neutrophils lies on a continuum due to their rapid aging, the effect of the gastrointestinal microbiome, and accrued cellular damage as neutrophils traverse narrow capillaries. Such phenotypic differences should not be regarded as neutrophil subpopulations if all neutrophils generated in the BM or PB can exhibit those features. Our analysis revealed five BM neutrophil subpopulations (including the GMP population) during development and maturation including three dividing subsets and two post-mitotic maturing subsets. The peripheral blood contained three distinct neutrophil subsets defined by distinct molecular signatures. Their identity was stable and largely maintained during bacterial infection.

The discrete and definable ISG-expressing G5b neutrophil subpopulation was present in both humans and mice, expanded during bacterial infection, and was independent of neutrophil activation. These cells highly expressed a set of ISGs and increased in number in *E. coli*-challenged hosts. Although G5b neutrophils were more similar to G5a than G5c neutrophils, the majority of G5b neutrophils directly developed from BM G4 cells (**Fig.7f**). In the spleens of *E. coli*-stimulated hosts, Gr1^+^ neutrophils were uniformly distributed in the red pup, while G5b cells preferentially localized to the subcapsular region, further demonstrating that the G5b neutrophil subpopulation is unique and suggesting that this subpopulation may be pre-programmed with a different host defense function. Interferon and interferon-related pathways are implicated in both viral and non-viral infections and play a critical role in host defenses (Schneider et al., 2014). ISG-related G5b neutrophils may be primed to combat invading pathogens even before infection occurs. Thus, the G5b neutrophil subset appears to be a functionally different population with distinct molecular signatures that maintain their features throughout the short neutrophil lifespan.

This study generated the first comprehensive single-cell RNA-seq map of neutrophils under physiological conditions and during bacterial infection. A number of neutrophil subpopulations have previously been identified through functional and phenotypic associations in various models, including but are not limited to human CD177^+^ (Hu et al., 2009; Silvestre-Roig et al., 2016; Wu et al., 2016; Zhou et al., 2018), olfactomedin-4^+^ (OLFM4) (Clemmensen et al., 2012), TCR-expressing (Puellmann et al., 2006), CD49d^+^VEGFR1^high^CXCR4^high^ angiogenic (Christoffersson et al., 2012; Massena et al., 2015), CD63^+^ (Tirouvanziam et al., 2008), IL-13^+^ (Chen et al., 2014), CD49^+^ (Cheung et al., 2010; Tsuda et al., 2004), IL-17-producing (Taylor et al., 2014), CD62L^dim^/CD16^bright^ and CD62L^bright^/CD16^dim^ (Pillay et al., 2012), immunosuppressive CD11c^bright^CD62L^dim^CD11b^bright^CD16^bright^ (Pillay et al., 2012), CD16^dim^ banded (Leliefeld et al., 2018), CD62L^dim^ (Tak et al., 2017), mature CD10^+^ and immature CD10^-^ (Marini et al., 2017), and tumor-elicited immature c-Kit^+^ (Rice et al., 2018) neutrophils. Additionally, Buckley et al. identified a subset of human CD54 (ICAM-1)^high^CXCR1^low^VEGFR^high^ neutrophils representing cells migrating through an endothelial monolayer and then re-emerging by reverse transmigration (RT) (Buckley et al., 2006). More recently, Deniset et al revealed that splenic Ly6G^high^ mature and Ly6G^int^ immature neutrophils contribute to eradication of *S. pneumoniae*. Resident neutrophil progenitors (CD11b^+^Ly6G^int^c-Kit^+^) in the spleen undergo emergency proliferation and mobilization from their splenic niche after pneumococcal stimulation to increase the effector mature neutrophil pool (Deniset et al., 2017). The molecular signatures of most of the aforementioned neutrophil subpopulations are unknown. Whether any of these neutrophil subpopulations overlap with or are derived from a particular neutrophil subset identified in current study will need to be determined.

Neutrophils are now known to be important in cancer. Two tumor-associated neutrophil (TAN) subpopulations are present in tumors: pro-inflammatory antitumorigenic N1 neutrophils and pro-tumorigenic N2 neutrophils (Fridlender et al., 2009; Giese et al., 2019; Sionov et al., 2015). N1 TANs showed a more immunostimulatory mRNA profile than N2 neutrophils, expressing higher levels of TNF-α, CCL3, iNOS, and ICAM-1. In contrast, N2 TANs express higher levels of CCL2, CCL5, CCL17, VEGF, and arginase, an important immunosuppressor of the adaptive immune system (Fridlender et al., 2009; Mishalian et al., 2013; Shaul et al., 2016). In a mouse melanoma model, IFN-β deficiency expands the number of TANs expressing elevated levels of genes encoding the proangiogenic factors VEGF and MMP9, CXCR4, and the transcription factors c-myc and STAT3, thereby augmenting tumor angiogenesis and growth (Jablonska et al., 2010). Sagiv et al. identified three distinct circulating neutrophil populations in cancer: high-density mature small anti-tumor neutrophils, low-density mature large pro-tumor neutrophils, and low-density immature large pro-tumor granulocyte-myeloid-derived suppressor cells (G-MDSC). MET expression, induced by inflammatory stimuli, is required for the recruitment of anti-tumor neutrophils (Finisguerra et al., 2015). Low-density neutrophils (LDN) were defined due to their appearance in the low-density layer of a Ficoll gradient (Carmona-Rivera and Kaplan, 2013; Garcia-Romo et al., 2011; Hacbarth and Kajdacsy-Balla, 1986). These cells may be responsible for sustaining chronic inflammation and have elevated capability to form NETs (Kanamaru et al., 2018; Khandpur et al., 2013). LDNs have a gene expression signature associated with a reduced inflammatory state (Sagiv et al., 2015). G-MDSC represents a group of immature neutrophils with immunosuppressive functions (Tcyganov et al., 2018; Veglia et al., 2018) characterized as CD11b^+^Ly6G^hi^Ly6C^lo^ in mice and CD14^−^CD11b^+^CD15^+^ (or CD66b^+^) in humans (Bronte et al., 2016). Condamine et al. identified lectin-type oxidized LDL receptor 1 (LOX-1) as a specific marker of human G-MDSCs associated with endoplasmic reticulum stress and lipid metabolism in cancer patients (Condamine et al., 2016). Comparative transcriptomic analyses of TANs, G-MDSCs, and naïve neutrophils suggested that these three subpopulations are distinct, with an unexpected close proximity of G-MDSCs and naïve neutrophils compared with TANs (Fridlender et al., 2012). Structural genes and genes related to cytotoxicity were significantly downregulated in TANs. In contrast, many immune-related genes and pathways, including genes related to the antigen-presenting complex (e.g., MHC-II complex genes) and cytokines (e.g., TNF-α, IL-1-α/β), were upregulated in G-MDSCs and further upregulated in TANs. We examined the expression of genes related to each tumor-related neutrophil subpopulation but could not correlate any of these neutrophil subsets with any of our novel subpopulations, suggesting that these specialized neutrophils are reprogrammed and acquire unique molecular signatures in the tumor-bearing host. Alternatively, these functionally defined neutrophil subsets, whose identities often rely on expression of a few surface markers, may still be heterogenous, leading to difficulty in recognizing the involved neutrophil populations. Interestingly, a group of ISG-expressing tumor-infiltrating neutrophils was recently identified in human and mouse lung cancers (Zilionis et al., 2019). Nevertheless, the transcriptome of this neutrophil population is different to that of G5b neutrophils, again indicating significant neutrophil reprogramming in the tumor microenvironment.

Using mass cytometry (CyTOF) and cell cycle-based analysis, Evrard et al. identified three neutrophil subsets within the bone marrow: committed c-Kit^low/int^ proliferative neutrophil precursors expressing primary and secondary granule proteins (preNeu), CXCR2^low^ non-proliferating immature neutrophils highly expressing secondary granule proteins, and CXCR2^high^ mature neutrophils highly expressing gelatinase granule proteins (Evrard et al., 2018). Based on these features, these three populations are likely correlated with G1/G2, G3, and G4 neutrophils identified in our study based on RNA-seq data (**Graphical Abstract**). More recently, a proliferative unipotent neutrophil precursor that suppresses T cell activation and promotes tumor growth was identified in the mouse bone marrow that generates neutrophils after intra-bone marrow adoptive transfer. ScRNA-seq analysis of these cells further revealed two populations: an early-stage ckit^+^Gfi1^low^Cebpa^hi^Ly6G^low^ progenitor with stem cell morphology and a late-stage ckit^+^Gfi1^hi^Cebpa^low^Ly6G^+^ precursor with morphological features similar to transient neutrophil precursors (Zhu et al., 2018). Further analysis suggested that the late-stage progenitors were mostly similar to the preNeu population identified by Evrard et al. (Zhu et al., 2018) (**Graphical Abstract**). Notably, in an earlier study, Kim et al. also defined a population of proliferative late-stage neutrophil precursors (NeuP) in the BM characterized by a lin^−^c-Kit^+^CD11b^+^Ly6G^lo^Ly6B^int^CD115^−^Gfi1^+^ signature (Kim et al., 2017) that should be located in the G1/G2 population. It was later found that this NeuP population was highly heterogeneous and contained other myeloid progenitors (G0 cells) (Zhu et al., 2018).

It is commonly believed that neutrophils mobilize from the BM to the PB only when they become fully mature. RNA velocity analysis revealed unexpected complexity of neutrophil mobilization from the BM to PB. Under homeostatic conditions, a significant number of G3 neutrophils were mobilized to the PB or tissue without going through the most mature G4 cells. Collectively, these results demonstrated that immature PB neutrophils in healthy hosts are mainly derived from more primitive early stage post-mitotic G3 cells, which were more mobile than the more mature G4 cells. Thus, full maturation may not be a pre-requisite for neutrophil mobilization. Early-stage post-mitotic G3 cells can be mobilized from the BM to the periphery via a maturation-independent mechanism to complete their development in the PB and/or SP (**Fig.7f**). Interestingly, the transformation from G3 to G5a cells was suppressed during infection and G3 cells in infected hosts predominantly differentiated to G4 cells.

Infection, long-term inflammation, and cancer can all reprogram granulopoiesis to produce neutrophils with different gene expression patterns. Indeed, we observed global changes in the neutrophil transcriptome in response to a bacterial stimulus. Nevertheless, the core identity of each neutrophil population and the overall neutrophil differentiation program remained the same during bacterial infection. The infection-triggered increase in neutrophils was mainly due to accelerated cell division in the more primitive G1 population rather than the G2 population, which was responsible for the majority of cell proliferation during homeostasis. One explanation for such a mechanism is that, under steady-state conditions, proliferation of G2 cells is already relatively high, so accelerating division of the more primitive G1 cells would provide a more efficient way to expand the neutrophil population. Infection also significantly shortened the period of post-mitotic maturation, leading to faster turnover of maturing neutrophils in the BM. Taken together, bacterial infection reprogrammed the structure of the neutrophil population and the dynamic transition between each subpopulation but did not produce distinct, functionally different neutrophil subpopulations in infected hosts.

## ACKNOWLEDGEMENTS

Cell sorting was performed at the HSCI/DRC Flow Core (NIH P30DK036836). H. Luo is supported by National Institutes of Health grants (1 R01 AI142642, 1 R01 AI145274, 1 R01 AI141386, R01HL092020, and P01 HL095489) and a grant from FAMRI (CIA 123008). Q.S. and C.L. were supported by Natural Science Foundation of China (31871266), Chinese National Key Projects of Research and Development (2016YFA0100103), and NSFC Key Research Grant 71532001. F. Ma is supported by the grants from National Basic Research Program of China (2015CB964903), Chinese Academy of Medical Sciences (CAMS) Innovation Fund for Medical Sciences (2017-I2M-1-015, 2016-I2M-1-017), the Non-profit Central Research Institute Fund of Chinese Academy of Medical Sciences (2018RC31002, 2018PT32034, 2017PT31033), National Natural Sciences Foundation of China (31271484 and 31471116), Natural Science Foundation of Tianjin City (18JCYBJC25700).

## AUTHOR CONTRI BUTIONS

Conceptualization, H.R.L., C.L., and F.M.; Methodology, H.R.L., L.S., and C.L.; Investigation, X.X., Q.S., P.W., X.Z., J.S., R.G., Q.R., S.Z., H.Y., S.P. and H.K.; Formal Analysis, X.X., Q.S., P.W., X.Z., J.S., R.G., Q.R., S.Z., H.Y., S.P. and H.K.; Writing – Original Draft, H.R.L., X.X., J.S., and Q.S.; Writing –Review & Editing, H.R.L., C.L., X.X., J.S., Q.S., and F.M.; Funding Acquisition, T.C., C.L., F.M., L.S. and H.R.L.; Resources, H.R.L., C.L., T.C., Y.X., and L.S.; Supervision, H.R.L., Y.X., C.L., and F.M.; Project Administration, H.R.L., C.L., Y.X., T.C., and F.M.

## STAR METHODS

### KEY RESOURCES TABLE

### LEAD CONTACT AND MATERIALS AVAILABILITY

Further information and requests for resources and reagents should be directed to and will be fulfilled by the Lead Contact, Hongbo R. Luo.

### EXPERIMENTAL MODEL AND SUBJECT DETAILS

#### Mouse strains

Female C57BL/6 mice were purchased from the Jackson Laboratory (Bar Harbor, ME). Eight-to-ten-week-old mice were used in all experiments. All animal experiments were conducted in accordance with the Animal Welfare Guidelines of the Children’s Hospital Boston. The Children’s Hospital Animal Care and Use Committee approved and monitored all procedures.

#### Mouse peritonitis model

Wild-type mice were intraperitoneally injected with 1×10^7^ *E. coli* (ATCC^®^ 19138™) in 300 μl PBS. At different timepoints after injection, mice were anesthetized with isoflurane, retro-orbital blood was collected, and then mice were sacrificed by euthanizing with CO2. Cells from different organs such as bone marrow, spleen, liver, and peritoneal exudate were collected as detailed below.

### METHOD DETAILS

#### Mouse neutrophil isolation

Peripheral blood was collected by retro-orbital bleeding. 600-800 ml peripheral blood was diluted with 3 ml HBSS containing 15 mM EDTA. Cells were centrifuged for 10 min at 500 x g. Red blood cells were lysed by resuspension in 5 ml ACK (ammonium-chloride-potassium) lysis buffer (Thermo Fisher Scientific, Waltham, MA) for 5 min at RT. 10 ml RPMI + 2% FBS were added to stop lysis followed by centrifugation at 500 x g for 5 min. Cells were washed twice with 10 ml HBSS + 2 mM EDTA + 1% BSA before being re-suspended in 500 μl PBS + 1% BSA. For bone marrow neutrophil isolation, whole bone marrow cells were flushed from the femur, tibia, and ilia leg bones with 5 ml HBSS + 2 mM EDTA + 1% BSA and filtered through a 70 μm cell strainer. Cells were centrifuged for 10 min at 500 x g. Red blood cells were lysed with 1 ml ACK lysis buffer for 2 min at room temperature (RT) and washed twice with HBSS + 2 mM EDTA + 1% BSA and re-suspended in 200 μl PBS + 1% BSA. c-kit-positive bone marrow cells were first enriched by positive selection using c-kit (CD117) microbeads (Miltenyi Biotec, Bergisch Gladbach, Germany) and further purified by FACS sorting c-kit-positive cells. To isolate spleen and liver neutrophils, spleens and livers were gently disaggregated through a 70 μm cell strainer with a 1 ml syringe plunger. Whole spleen and liver cells were collected and red blood cells lysed using the same procedure as bone marrow cells. Finally, peritoneal cavity exudate cells were harvested by three successive washes with 10 ml HBSS + 15 mM EDTA + 1% BSA. After centrifugation, they were washed twice with the same solution and the cells re-suspended in 100 μl PBS + 1% BSA. Of note, neutrophils display circadian oscillations in number and phenotype, and neutrophil aging is an intrinsically-driven *bona fide* circadian process (Adrover et al., 2019). Thus, all samples in this study were prepared from mice sacrificed at the same time each morning.

#### Human sample collection

Peripheral blood was collected from a male healthy donor aged 32 years in a heparin anticoagulant tube. 10 ml (100% of blood volume) of 6% hydroxyethyl solution was added into the heparinized blood and inverted gently several times for adequate mixing. The blood was kept at RT for 30-45 min before pipetting the supernatant into a 50 ml Falcon tube followed by centrifugation at 290 x g for 5 mins without braking. Cells were washed twice and lysed with ACK to completely remove red blood cells. Samples were stained with Percp-cy5.5-conjugated anti-human CD33 antibody for 20 min and DAPI was added to cells prior to sorting by FACS with a FACSAria III cell sorter (BD Biosciences, Franklin Lakes, NJ). The Ethics Committee of Tianjin Blood Disease Hospital approved the study protocol, and the donor provided written informed consent for sample collection and data analysis.

#### Single cell collection, library construction, and sequencing

Single cell suspensions were stained for 30 min at 4°C with fluorophore-conjugated antibodies (APC/CY7-conjugated anti-Gr1, FITC-conjugated-anti-CD45), filtered through 40 μm cell strainers, and DAPI was added prior to sorting by FACS with a FACSAria III cell sorter (BD Biosciences). Designated cells were sorted into PBS containing 0.05% BSA following the 10X Genomics protocol. Cell preparation time before loading onto the 10X Chromium controller was <2 h. Cell viability and counting were evaluated with trypan blue by microscopy, and samples with viabilities >85% were used for sequencing. Libraries were constructed using the Single Cell 3’ Library Kit V2 (10X Genomics, Pleasanton, CA). Transcriptome profiles of individual cells were determined by 10X genomic-based droplet sequencing. Once prepared, indexed cDNA libraries were sequenced with paired-end reads on an Illumina NovaSeq6000 (Illumina, San Diego, CA).

#### Bulk RNA isolation and sequencing

BM cells were prepared as previously described. BM cells were first stained with: biotin-conjugated anti-CD4; biotin-conjugated anti-CD8a; biotin-conjugated anti-Ter119; and biotin-conjugated anti-B220/CD45R antibodies for 20 min and then stained with PE/cy7-conjugated streptavidin; APC-conjugated anti-c-kit; PE-conjugated anti-Ly6G; FITC-conjugated anti-CD34 antibodies for 90 min at 4°C. MB, PM, MM, MC, mature band, and segmented neutrophils were sorted with a Moflo cell sorter (Beckman Coulter, Brea, CA). Total RNA was extracted from those populations using the Qiagen RNeasy Mini Kit (Qiagen, Hilden, Germany). RNA quality was evaluated spectrophotometrically, and the quality was assessed with the Agilent 2100 Bioanalyzer (Agilent Technologies, Santa Clara, CA). All samples showed RNA integrity >7.5; RNA-seq libraries were prepared using the KAPA mRNA HyperPrep Kit (Illumina). Once prepared, indexed cDNA libraries were pooled in equimolar amounts and sequenced with paired-end reads on an Illumina HiSeq2500.

#### Wright-Giemsa staining and examination of morphology-defined neutrophil populations

The first recognizable cells of neutrophil lineage in the BM are myeloblasts (MBs), which are characterized by a high nuclear-to-cytoplasmic (NC) ratio and dispersed chromatin. MBs then irreversibly differentiate into promyelocytes (PMs), which are characterized by a round nucleus and azurophil granules, followed by myelocytes (MCs) characterized by a round nucleus and specific granules. Metamyelocytes (MMs) are characterized by nuclear indentations (kidney-shaped nuclei) and the emergence of secretary vesicles. Finally, MMs are divided into band cells (BCs) with a band-shaped nucleus and segmented cells (SCs, aka polymorphonuclear granulocytes) with a segmented nucleus. Cells were sorted and concentrated onto microscope slides by cytospinning. Slides were dried and stained using the Diff-Quick Stain Set (Siemens, Munich, Germany). Stained slides were rinsed under running tap water and air-dried for 10 min. Images were obtained under a microscope with a 63x objective.

#### EDU incorporation assay

5-ethynyl-2’-deoxyuridine (EdU), a thymidine analogue, can track cells post-mitotically in the BM and PB (**Fig.6e**). EdU is incorporated into DNA in the S-phase of the cell cycle, and the half-life of EdU is only about 30 min (Cheraghali et al., 1995), so incorporation can only occur in the first 1-2 h after EdU intraperitoneal injection. After 1 h of i.p. injection with 0.5 mg EdU, mice were injected with *E. coli* as above to induced peritonitis. Mice were sacrificed at designated timepoints, and BM, blood, and spleen cells were harvested followed by staining with fluorescent-conjugated antibodies: APC-conjugated anti-CD11b; APC/cy7-conjugated anti-Ly6g; and PE-conjugated anti-CXCR4 antibodies. Labeled cells were fixed, permeabilized, and stained with azide dye using an EdU Proliferation Kit (BD Biosciences). Cells were further washed and analyzed using a BD FACSCanto II (BD Biosciences). Data were analyzed using FlowJo software (FloJo, BD Biosciences).

#### Spleen cryosection preparation

Spleens were fixed in 1% formaldehyde (StatLab, McKinney, TX) for 4–8 h, rehydrated in 30% sucrose solution for 72 h, and snap frozen in OCT (Sakura Finetek, Japan). Single-cell-thick (5 μm) spleen cryosections were obtained using a Leica Cryostat and the CryoJane tape transfer system (Leica Microsystems, Wetzlar, Germany). For immunofluorescent staining, slides were rehydrated in PBS for 10 min followed by rinsing in PBST (PBS + 0.1% Tween20); blocking was performed with PBS + 10% donkey serum for 20 min; the diluted primary rat anti-S100a8 (Thermo Fisher Scientific #335806) and rabbit anti-Ifit1 (Abcam, Cambridge, UK #ab236256) antibodies were added and incubated for 1 h at RT. After 3x washes with PBST, AlexaFluor 488-conjugated donkey anti-rat antibody (Jackson ImmunoResearch, West Grove, PA #141697) and Cy^TM^3-conjugated donkey anti-rabbit antibody (Jackson ImmunoResearch #143460) were added and incubated for 30 min at RT. Slides were washed 3x with PBST and then stained with DAPI (0.5 μM) for 3 min. Slides were rinsed in PBS and were covered with mounting solution (Vectashield, Vector Laboratories, Burlingame, CA).

#### Laser scanning cytometry (LSC)

Laser scanning cytometer (LSC) is an emerging technology that images and quantitatively analyzes cellular and subcellular criteria within tissues, re-interrogating identified cell subpopulation(s) for *in situ* characterization of the molecular and cellular events associated with those cells (Harnett, 2007; Kwak et al., 2015). LSC was performed with an iCys Research Imaging Cytometer four-laser system (Thorlabs, Newton, NJ). Each section was first scanned with a 10x objective using the 405 nm laser to generate low-resolution images of the DAPI-stained nuclei and obtain a general view of the spleen. Subsequently, the sections were divided into small regions and scanned with a 40x dry objective lens to create high-resolution field images. Data were analyzed using iCys Cytometric Analysis Software (Thorlabs).

#### Confocal imaging

20 μm thick sections were prepared and stained as described above. Confocal images were obtained using the Zeiss LSM 700 Laser Scanning Confocal microscope (Carl Zeiss AG, Oberkochen, Germany). Data were analyzed using Imaris Software (Oxford Instruments, Abingdon, UK).

### QUANTIFICATION AND STATISTICAL ANALYSIS

#### scRNA-seq data processing

The quality of sequencing reads was evaluated using FastQC and MultiQC. Cell Ranger v2.2.0 was used to align the sequencing reads (fastq) to the mm10 mouse transcriptome and quantify the expression of transcripts in each cell. This pipeline resulted in a gene expression matrix for each sample, which records the number of UMIs for each gene associated with each cell barcode. For human data, sequenced reads were aligned to the hg38 human transcriptome, then quantify the expression of transcripts in each cell using BD™ Rhapsody Whole Transcriptome Assay Analysis Pipeline. Unless otherwise stated, all downstream analyses were implemented using R v3.5.2 and the package Seurat v2.3.4 (Butler et al., 2018). Due to dissimilar data qualities, low-quality cells were filtered using sample-specific cutoffs (Table S1). The NormalizeData function was performed using default parameters to remove the differences in sequencing depth across cells.

For experiment described in Fig 1, cells from four samples were pooled and analyzed together. After rigorous quality control, we obtain 19,582 high-quality cells with average 1,241 genes per cell profiled, resulting in a total of 18,269 mouse genes detected in all cells (Fig.S1f). Clusters G1 to G5 were neutrophils at different maturation stages. G1 and G2 were early-stage neutrophils with higher expression of *Elane*, *Mpo*, *Fcnb*, and *Camp* (Fig.1d-e). Neutrophils are terminally differentiated. The transition from a proliferative cell to terminal differentiation was accompanied by a dramatic change in expression of important cell-cycle regulatory proteins, so we next performed a single-cell resolution analysis of cell cycle activation during neutrophil differentiation based on the expression of G1/S and G2/M phase-specific genes (Kowalczyk et al., 2015; Tirosh et al., 2016b) (Fig.1i). Cells in the G0 to G2 stages underwent active proliferation, while cell division stopped abruptly thereafter. CDC28 Protein Kinase Regulatory Subunit 2 (CKS2), Mki67, and Cdc20 were all strongly downregulated at the mRNA level.

For experiment described in Fig 4, After excluding low-quality cells, a total of 25,897 cells, including 4421 cells from the BM (eBM_Gr1), 6232 cells from the PB (ePB_Gr1), 5989 cells from the SP (eSP_Gr1), 4435 cells from the liver (eLV_Gr1), and 4169 cells from the peritoneal cavity (ePC_Gr1) were available for analysis (Fig.S4b-c).

#### Batch correction

There was substantial variability between cells obtained from different samples, likely reflecting a combination biological and technical differences. In this case, the batch had little effect on partitioning cell types and thus cell clustering into neutrophils, B cells, T cells, monocytes, dendritic cells, erythrocytes, and progenitors.

However, when clustering neutrophils alone, cells clustered first by sample rather than by biological clusters. Therefore, the ScaleData function was used to eliminate cell-cell variation in gene expression driven by batch and mitochondrial gene expression. Importantly, the batch-corrected data were only used for principal component analysis (PCA) and all steps relying on PCA (e.g., clustering, UMAP visualization); all other analyses (e.g., differential expression analysis) were based on the normalized data without batch correction.

#### Dimension reduction

Dimension reduction was performed at three stages of the analysis: the selection of variable genes, PCA, and uniform manifold approximation and projection (UMAP) (Becht et al., 2018b). The FindVariableGenes function (y.cutoff = 1 for control total cells; y.cutoff = 1.2 for control neutrophils; y.cutoff = 0.7 for *E. coli*-challenged total cells) was applied to select highly variable genes covering most biological information contained in the whole transcriptome. Then, the variable genes were used for PCA implemented with the RunPCA function. Next, we selected PCs 1-20 (for total cells) or 1-15 (for neutrophils) as input to perform the RunUMAP function to obtain bidimensional coordinates for each cell.

#### Unsupervised clustering

We performed the FindClusters function (resolution 0.3, 0.6, and 0.2 for control total cell, neutrophils, and *E. coli*-challenged total cells, respectively) to cluster cells using the Louvain algorithm based on the same PCs as RunUMAP function.

#### Identification of differentially expressed genes

We used the FindMarkers or FindAllMarkers function (test.use = ‘‘t’’, logfc.threshold = log(2)) based on normalized data to identify differentially expressed genes (DEGs). P-value adjustment was performed using Bonferroni correction based on the total number of genes in the dataset. DEGs with adjusted p-values>0.05 were filtered out. Gene ontology (GO) analysis was performed by using the R package clusterProfiler (Yu et al., 2012). In experiment described in Fig S5, we conducted differential gene expression analysis in each neutrophil subpopulation using the non-parametric Wilcoxon rank sum test and identified DEGs with an average expression fold-change >2.

#### Developmental trajectory inference

Pseudotime was generated with Monocle v2 (Qiu et al., 2017b) to infer the potential lineage differentiation trajectory. The newCellDataSet function (lowerDetectionLimit = 0.5, expressionFamily = negbinomial.size) was used to build the object based on the above highly variable genes identified by Seurat v2.3.4.

#### Bulk RNA-seq analysis

The quality of sequencing reads was evaluated using FastQC and MultiQC. Adaptor sequences and low-quality score bases were trimmed using trimmomatic/0.36. The resulting reads were then mapped to the mouse reference sequence (GRCm38/mm10, Ensemble release 81) and counted using STAR2.5.2b alignment software. Gene differential expression analysis was performed using the R package EdgeR.

#### Scoring of biological processes

Individual cells were scored for their expression of gene signatures representing certain biological functions. For all signatures except neutrophil aging, functional scores were defined as the average normalized expression of corresponding genes. Aging score was defined as the weighted average of Z-scores of age-related genes, where the Z-scores were calculated by scaling the normalized expression of a gene across all cells. Gene weights were set to either 1 or −1 to reflect positive or negative relationships. The neutrophil maturation signature was derived by identifying the top 50 DEGs with highest fold changes and adjusted p-values <0.05 between mature cluster (G4) with immature clusters (G0-G3). Granule signatures were from (Cowland and Borregaard, 2016b). Other functional signatures were derived from the GO database (Consortium, 2018), with the full gene list provided in Table S4. For instance, to access the phagocytosis function at the transcript level, we determined a “phagocytosis score” by calculating the average expression of genes in the GO term “phagocytosis, engulfment” (GO:0006911). Apoptosis score was measured by the upregulation of the integrated proapoptotic pathway and downregulation of pro-survival gene expression (Fig 3b). To further dissect apoptotic heterogeneity in G5 populations independently of transcriptome-based sub-clustering, we fit a two-component Gaussian mixture model to the apoptotic score of all G5 cells using the R package mixtools version 1.1.0 (Benaglia et al., 2009). We then chose the distribution with the higher mean as the apoptotic group and assigned each cell to one of the two groups based on its posterior (Fig 3c).

Age-related genes were summarized from the previous literature (**Fig 2i**). Related to function, aged neutrophils express less adhesion molecule L-selectin (Cd62L, Sell) and more CD11b (αM, Itgam), lymphocyte function-associated antigen-1 (CD11a/β2), CD49d (integrin α4, Itga4), TLR4, ICAM-1, CD44, and CD11c (Itgax) (Casanova-Acebes et al., 2013; Uhl et al., 2016; Zhang et al., 2015). Additionally, aged neutrophils express more surface CXCR4 and less CXCR2, which regulates their release from and return to the BM (Eash et al., 2010; Eash et al., 2009; Martin et al., 2003; Nagase et al., 2002; Zhang et al., 2015). CXCR4 may also play a role in clearing aged, senescent neutrophils, particularly at BM sites. Anti-CXCR4 antibodies or CXCR4 antagonists impede neutrophil homing to the BM (Martin et al., 2003; Suratt et al., 2004). Finally, aged neutrophils exhibit increased expression of CD24, a GPI-linked glycoprotein which induces apoptosis when crosslinked (Parlato et al., 2014) and reduced expression of CD47, the “don’t eat me” signal that inhibits efferocytosis, a process leading to clearance of dead neutrophils (Jaiswal et al., 2009; Zhang et al., 2015).

ROS-mediated pathogen killing is a major host defense mechanism. In neutrophils, ROS are mainly produced by the phagocytic NADPH oxidase (aka the NOX2 complex). During cell activation, cytosolic components of the NADPH oxidase NCF2 (p67phox), Rac1 and/or Rac2, NCF4 (p40-phox), and NCF1 (p47phox) are recruited to the membrane to form a complex with membrane proteins CYBA (p22-phox) and CYBB (gp91 or cytochrome-b 558 subunit beta) (Babior et al., 2002; Dahlgren and Karlsson, 1999; Henderson and Chappel, 1996; Heyworth et al., 2003; Luo and Loison, 2008; Nauseef and Borregaard, 2014; Segal et al., 2000; Subramanian and Luo, 2009). We evaluated the “NADPH oxidase score” based on the expression of the seven NADPH oxidase-related genes (Fig S3d)

#### Comparison of scRNA-seq-defined populations with morphology-defined neutrophil subpopulations

To benchmark single-cell transcriptomic neutrophil classification against existing morphological classification schemes, we deconvoluted bulk RNA-seq profiles based on expression of scRNA-seq-identified group-specific signatures. This approach was similar to other existing deconvolution methods like CIBERSORT (Newman et al., 2015), but we used a linear regression model with the constraint of non-negative coefficients (i.e., non-negative least-squares problem) instead of the linear support vector regression in CIBERSORT. Although we manually chose 20 genes with highest fold-changes as signatures for each single cell group, we noted that the deconvolution in our case was robust to the choice of signatures. The regression model was built using R package nnls version 1.4 (Katharine M. Mullen, 2012). Bulk profiles were quantile normalized.

At different morphology-defined neutrophil differentiation stages, neutrophils produce different granules containing distinct enzymes and antimicrobial compounds. Thus, we also examined the expression of various granule genes in differentiating neutrophils. Genes related to primary (azurophilic) granules such as *Mpo* started to be expressed in some G0 cells, peaked in G1 cells, and then rapidly decreased in G2 cells (Fig.2a-b). MPO-negative granules can be divided into granules containing lactoferrin (LTF) but no gelatinase (MMP9), granules that contain both, and granules that contain gelatinase but no lactoferrin (Kjeldsen et al., 1993). We found sequential production of these granules in maturing neutrophils, with lactoferrin-containing granules emerging in G2 cells, lactoferrin and gelatinase-containing granules emerging in G3 cells, and gelatinase-containing granules (Ltf low) emerging in G4 cells (Fig.2a-c). Of the proteins that localize exclusively to secretory vesicles such as Frp1 and Vamp2, their cognate mRNA profiles peaked in G4 cells in the BM and continued to be expressed in PB neutrophils.

#### SCENIC analysis

SCENIC is a computational tool that infers regulatory modules or regulons by analyzing co-expression of transcription factors (TFs) and their putative target genes characterized by enrichment of corresponding TF-binding motifs in their regulatory regions (Aibar et al., 2017). Regulatory network analysis was performed on all control and *E.coli-*challenged samples using Python package pySCENIC version 0.9.11 (Aibar et al., 2017) with default parameters. We scaled the network inference step by first inferring regulons on a 6000-cell subset, then calculated AUCell scores for all 32,888 cells included in this analysis. Specifically, we randomly sampled 300 cells from each neutrophil population in each condition and 1200 non-neutrophil cells as the training set for network inference. Output co-expression modules were trimmed with cisTarget databases (mm10_refseq-r80_500bp_up_and_100bp_down_tss, mm9-tss-centered-10kb-7species, and mm9-500bp-upstream-10species). The identified 413 regulons were then scored to determine their activities in each cell. K-means clustering was performed on the first 20 principle components (PCs) of regulon activity matrix with cluster number k = 7.

#### Differential activity analysis of SCENIC regulons

To assess the effect of biological conditions on regulon activity, we applied a generalized linear model (GLM) as reported in (Lambrechts et al., 2018). We compared the AUCell score of each regulon with different baseline clusters corresponding to different biological questions like neutrophil cluster transition and infection response. GLM results were further filtered by p-values and visualized using R package ComplexHeatmap (Gu et al., 2016).

#### RNA velocity analysis

Cell RNA velocity analysis was performed using Velocyto program (La Manno et al., 2018a). This approach uses the relative proportion of unspliced and spliced mRNA abundance as an indicator of the future cell state (La Manno et al., 2018b). The calculated RNA velocity is a vector that predicts individual cell transition, with the direction and speed of each transition assessed based on the amplitude and direction of individual cell velocity arrows on the UMAP plot. Accordingly, the hierarchical relationship between two cell populations can be inferred by the directional flow in the RNA velocity vector field. Annotation of spliced and unspliced reads was first performed using velocyto.py command-line tools. Then, downstream analysis was performed using the velocyto.R pipeline. We retained the genes expressed in at least one cell population. In total, 4815 genes were used for the analysis. RNA velocities of each cell were estimated using the gene.relative.velocity.estimates function. Finally, the velocity field was projected onto the existing UMAP space.

#### Cell label transfer

Total cells were partitioned into distinct cell types annotated by the expression of known marker genes. Neutrophils in their steady state were partitioned into eight clusters based on gene expression profiles annotated according to their development order. *E. coli*-challenged neutrophils were annotated using a well-accepted method (Stuart et al., 2019). We first identified pairwise correspondences (aka anchors) between single cells across datasets (before and after *E. coli* challenge) to quantify the batch effect. Each cell in the *E. coli*-challenged dataset was then annotated based on the transcriptomic similarity between this cell and cells in the reference dataset. Specifically, cells would receive corresponding labels with the highest similarity scores, whereas cells with the highest similarity score lower than 0.5 were defined as “unassigned” and were not represented in the reference. In this way, each neutrophil from the new stimulated dataset was assigned a cluster name, and neutrophils sharing similar transcriptomic profiles would be placed into the same cluster. Hence, each cell in the bacterial infection state was assigned to one of the nine cluster labels. This transfer procedure was implemented using the FindTransferAnchors (dims = 1:15) and TransferData (dims = 1:15) functions in Seurat v3.0.2 (Stuart et al., 2019) with the combination of top 100 DEGs of each cluster.

### DATA AND CODE AVAILABILITY

All sequencing data have been deposited at NCBI GEO depository and are accessible with the accession number GSEXXX.

## Supplementary materials

**Figure S1.**
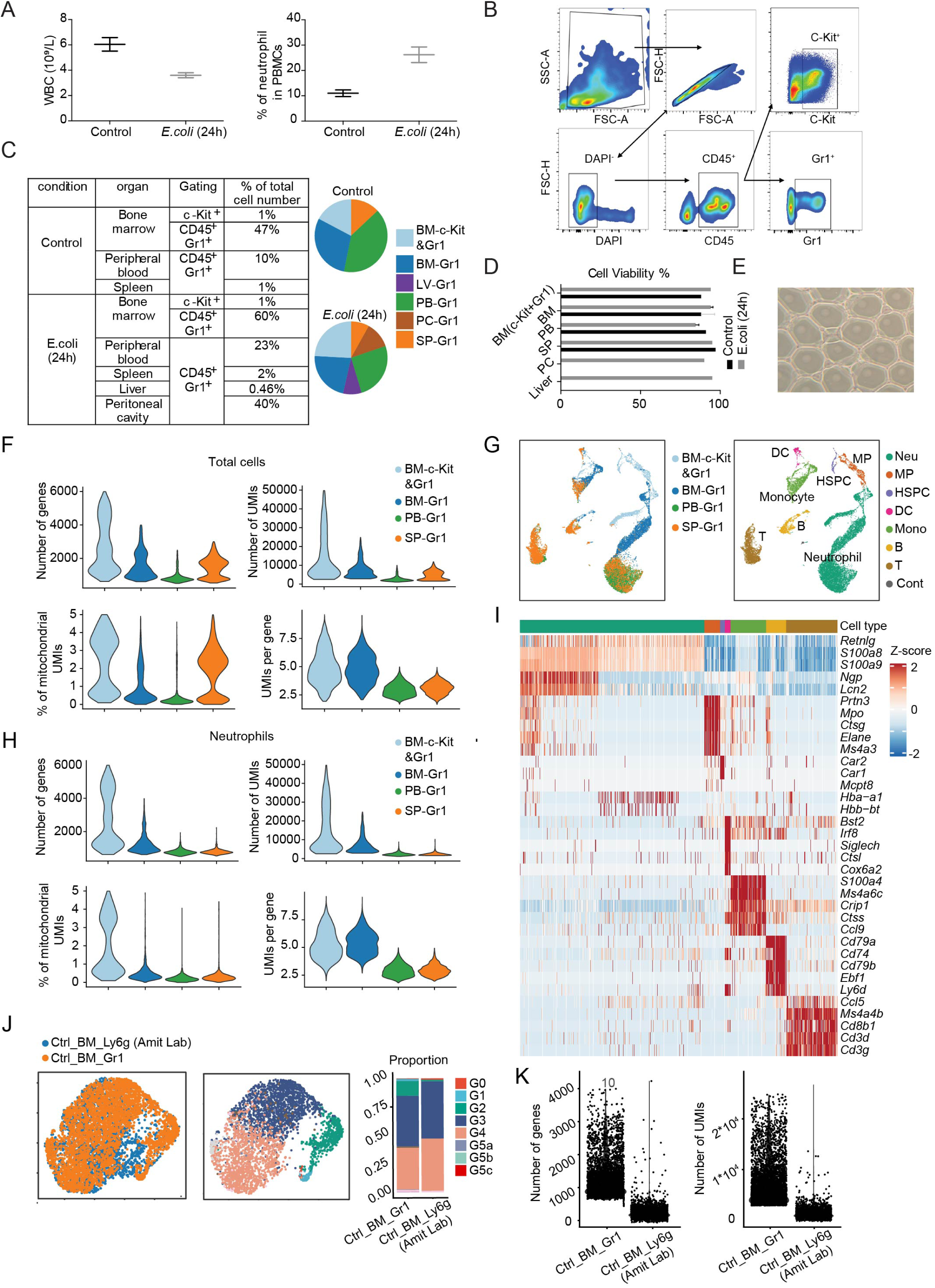
Sample preparation, quality controls, and related parameters and results related to scRNA-seq analysis. (A) Number of white blood cells and the proportion of neutrophils in mice before and after *E. coli* challenge evaluated by a hematology analyzer (Mindray BC-5000 Vet). Results are the mean ± SD of three independent experiments. (B) Fluorescence-activated cell sorting (FACS) strategy for scRNA-seq sample preparation. (C) Summary of sample information. Organ distributions of neutrophils before and after *E. coli* challenge are shown on the right. (D) Cell viability percentages immediately before cells were loaded into the 10X Chromium Controller. (E) Representative GEM formation after the 10X Chromium Controller under the microscope. (F) Violin plots of the number of genes, number of UMIs, mitochondria count percentage, and UMI per gene of all QC-passed cells in different organs. (G) Uniform manifold approximation and projection (UMAP) of 19,582 cells from the bone marrow (BM), peripheral blood (PB), and spleen (SP) colored by sample origin and cell type, respectively. Expression of unique genes specifically distinguished each cluster and associated them with neutrophils (Neu) (*S100a8* and *S100a9*), myeloid progenitors (MP) (*Cd34, Kit, Mpo* and *Elane*), hematopoietic stem progenitor cells (HSPC, not including MP) (*Cd34, Kit, Mpo^-^* and *Elane^-^*), monocytes (Mono) (*S100a4* and *Ccl9*/MIP-1γ), B cells (*Cd79a* and *Cd79b*), T cells (*CD3d* and *Ccl5*), and dendritic cells (DC) (*Siglech*), respectively. Cont: contaminated cells. (H) As in (F) but using only neutrophils in different organs. (I) Heatmap showing the five highest differentially expressed genes (DEGs) per cell type for all QC-passed cells. (J) Comparison of *Gr1^+^* BM neutrophil populations in our data with *Ly6g^+^* BM neutrophil populations in Dr. Ido Amit’s data. Cluster labels are transferred from our data to Dr. Ido Amit’s data (Giladi et al., 2018)(Methods). Left: UMAPs of 3591 *Gr1^+^* neutrophils and 2304 *Ly6g^+^* neutrophils colored by data set or cluster identity. Right: Neutrophil compositions in our data and Dr. Ido Amit’s data. (K) Violin plots of the number of genes and number of UMIs of our *Gr1^+^* neutrophils and Dr. Ido Amit’s *Ly6g^+^* neutrophils.

**Figure S2.**
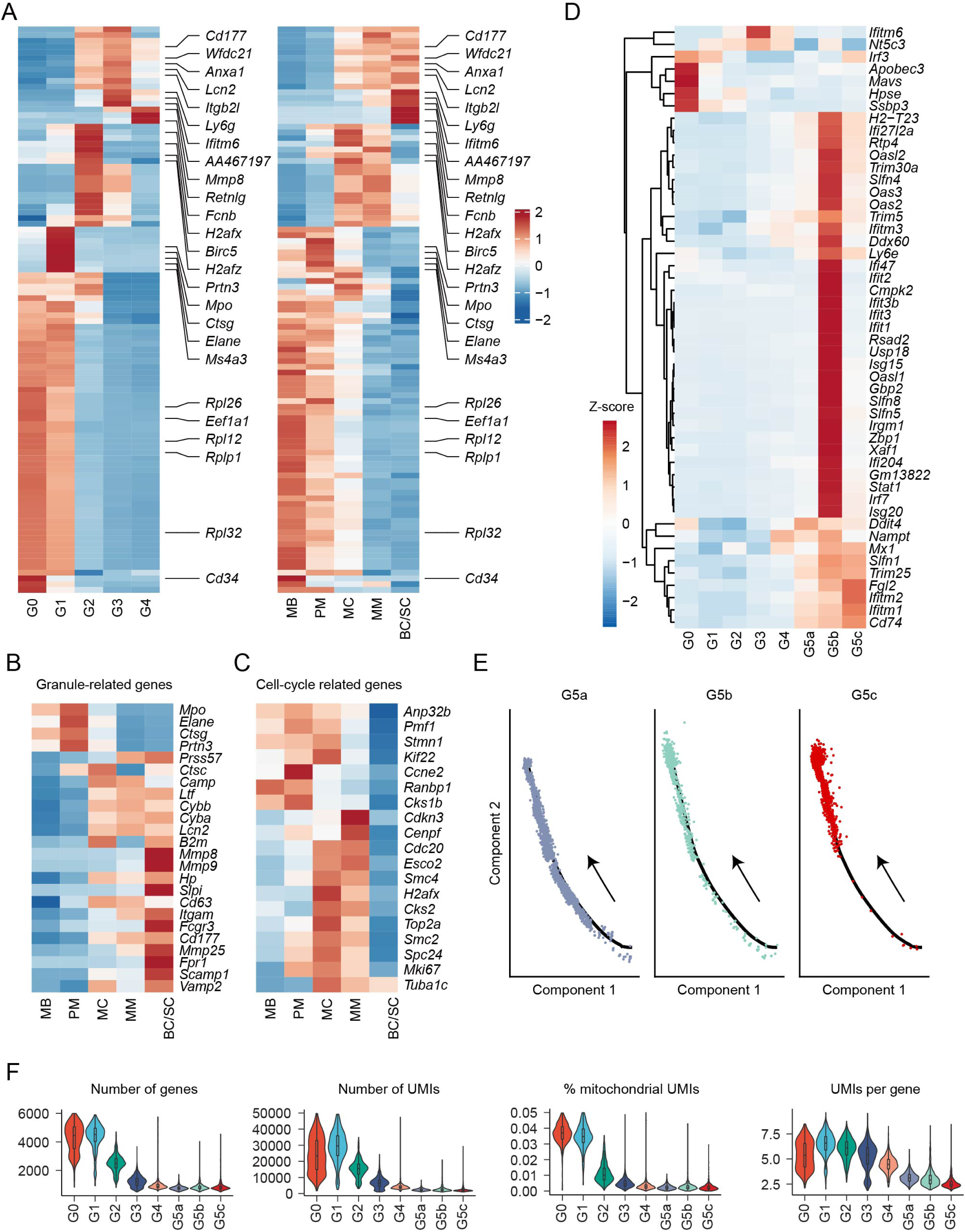
(A-C) scRNA-seq defined differentiating neutrophil populations correlated with classical morphology-defined neutrophil subpopulations. (A) Heatmaps showing row-scaled expression of the 20 highest DEGs per cluster across averaged single-cell groups (left) and morphological groups (right). Only genes detected in both scRNA-seq data and bulk RNA-seq data are visualized. Representative genes are indicated. (B-C) Heatmaps showing row-scaled expression of granule-related genes (B) and cell-cycle related genes (C) for morphological groups. **(D)** Heatmap showing row-scaled expression of 47 interferon-stimulated genes for each averaged cluster. **(E)** Monocle trajectories of neutrophil population G5a, G5b, and G5c. Each dot represents a single cell. **(F)** Violin plots of the number of genes, number of UMIs, mitochondria count percentage, and UMI per gene of neutrophils in each cluster.

**Figure S3.**
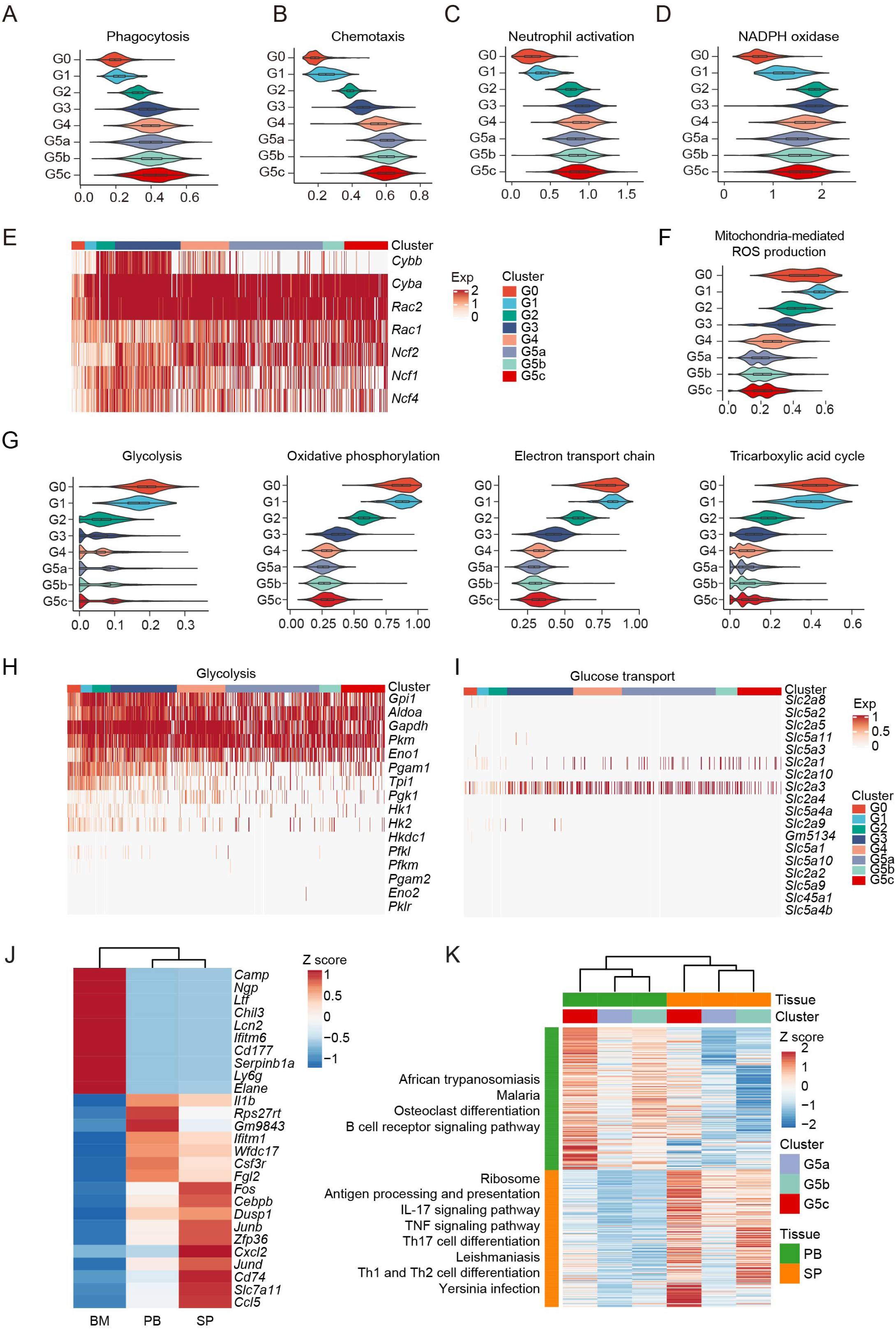
**(A-I) Characterization of neutrophil subpopulations.** (A-D) Violin plot of phagocytosis score (GO:0006911), chemotaxis score (GO:0030593), neutrophil activation score (GO:0042119), and NADPH oxidase score for each cluster. (E) Heatmap showing relative expression of seven genes of the NADPH oxidase complex for all neutrophils. (F) As in (A-D) but displaying mitochondria-mediated ROS production score (reactive oxygen species biosynthetic process, GO:1903409) for each cluster. (G) Violin plots of metabolic scores for each cluster. Glycolysis (Reactome Pathway Database #R-MMU-70171); Oxidative phosphorylation (GO:000619); Electron transport chain (GO:0022900); Tricarboxylic acid cycle (GO:0006099). (H-I) Heatmaps showing relative expression of glycolysis-related genes and glucose transport-related genes. **(J-K) Organ-specific transcriptome features.** (J) Heatmap showing row-scaled expression of the ten highest DEGs per organ for each averaged organ profile. (K) As in (J) but for each G5 subpopulation between PB and SP. KEGG analysis of DEGs for each G5 in these two organs. Left: selected KEGG terms with Benjamini-Hochberg-corrected P-values < 0.05 are shown.

**Figure S4.**
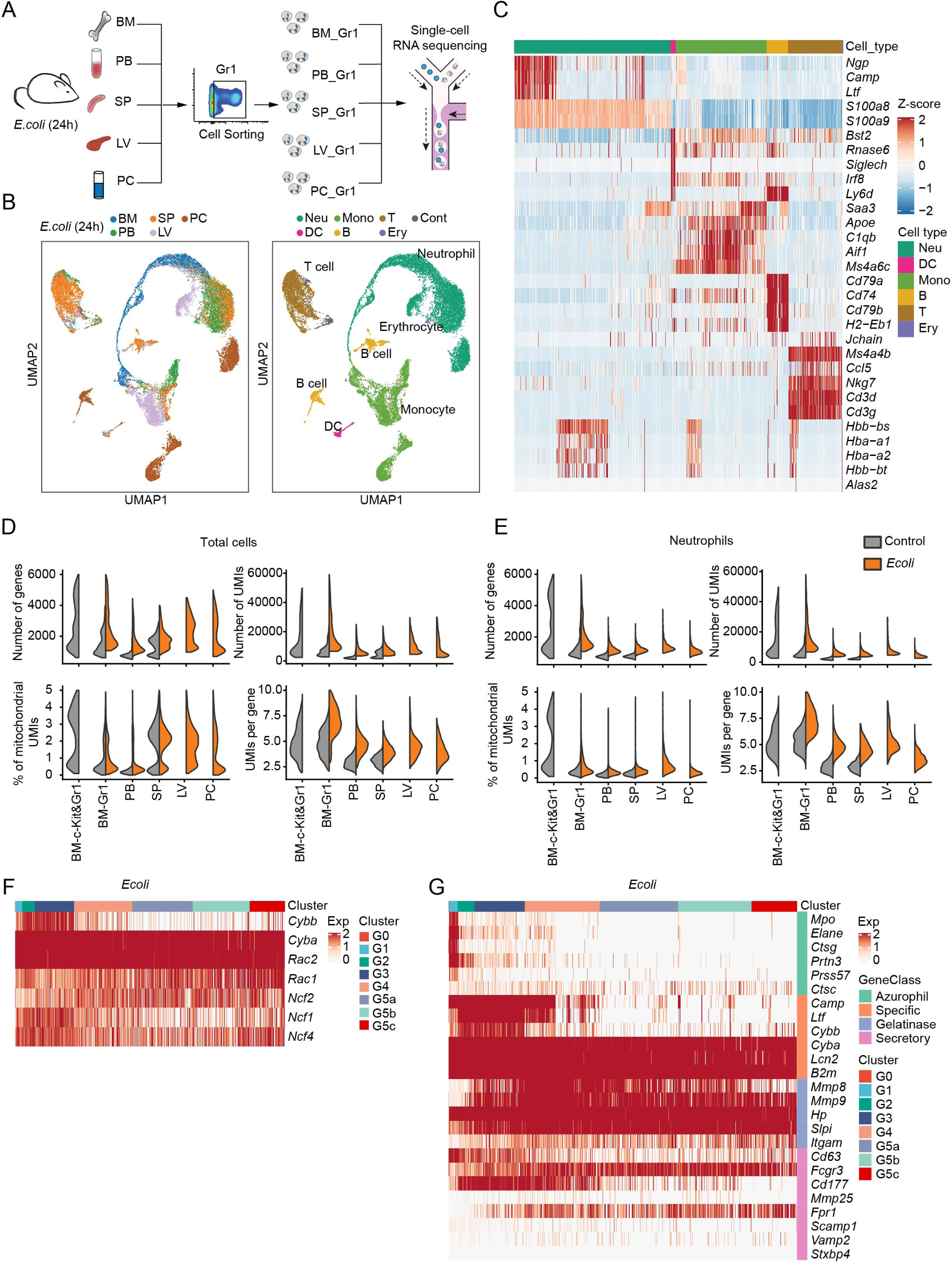
Single cell RNA-seq analysis of neutrophils in *E. coli*-challenged mice. (A) Experimental scheme of the sample collection process after *E. coli* challenge. (B) UMAPs of all 24,943 cells from BM, PB, and SP from *E. coli*-challenged mice colored by sample origin and cell type, respectively. (C) Heatmap showing row-scaled expression of the five highest DEGs for all QC-passed cells colored by cell type. (D) Comparisons of the number of genes, number of UMIs, mitochondria count percentage, and UMI per gene of all QC-passed cells in each organ before and after *E. coli* challenge. (E) As in (D) but only of all neutrophils in each organ. (F-G) Heatmaps showing expression of 7 genes of the NADPH oxidase complex (F) and neutrophil granule-related genes (G) for all neutrophils.

**Figure S5.**
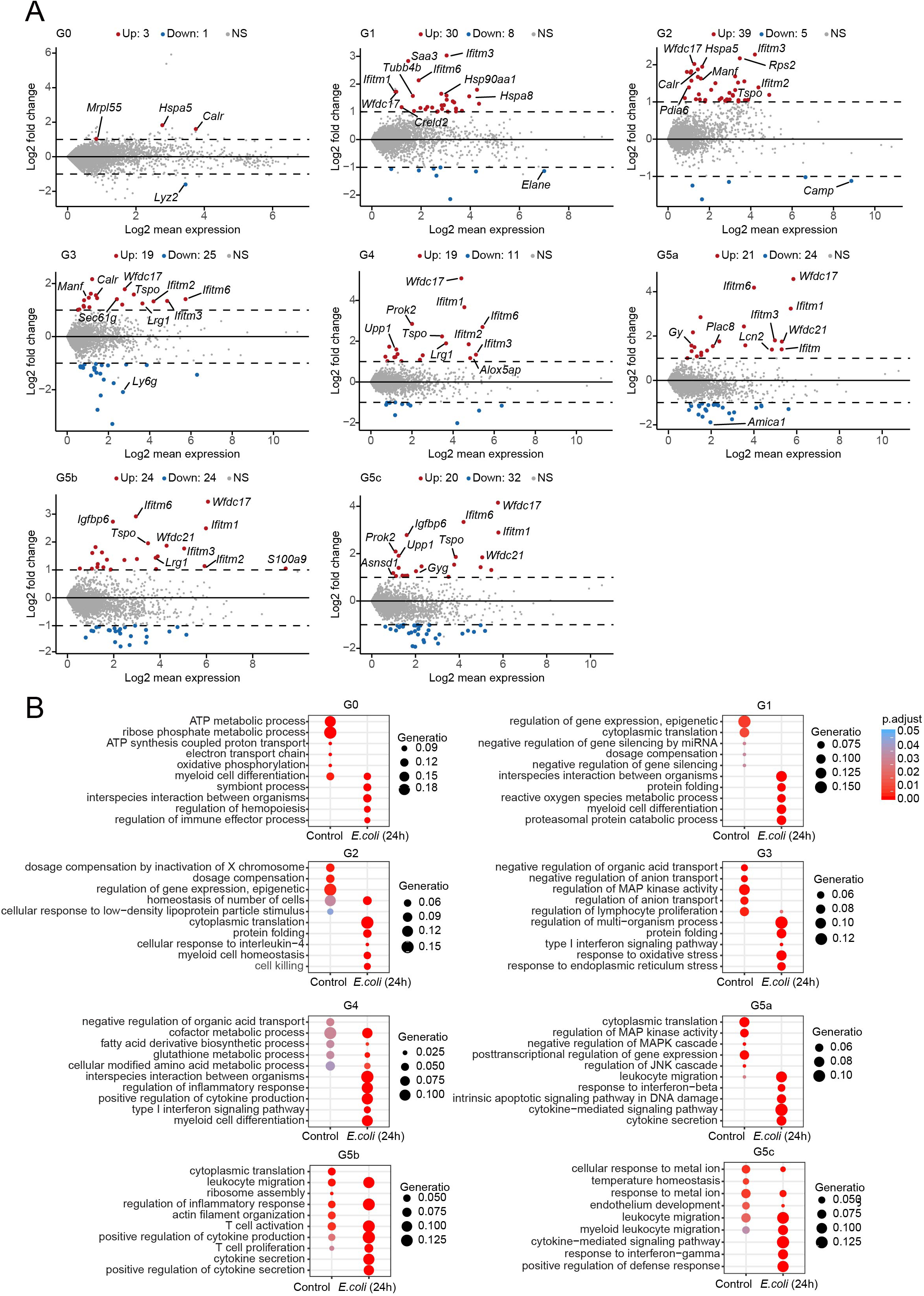
Differentially expressed genes in each neutrophil subpopulation in *E. coli*-challenged mice. (A) MA plots displaying genes that are up- (red) or downregulated (blue) after *E. coli* challenge for each cluster. Dashed lines denote fold change thresholds used when identifying DEGs. (B) Gene ontology (GO) analysis of DEGs before and after *E. coli* challenge for each cluster. Selected GO terms with Benjamini-Hochberg-corrected P-values < 0.05 (one-sided Fisher’s exact test) are shown.

**Figure S6.**
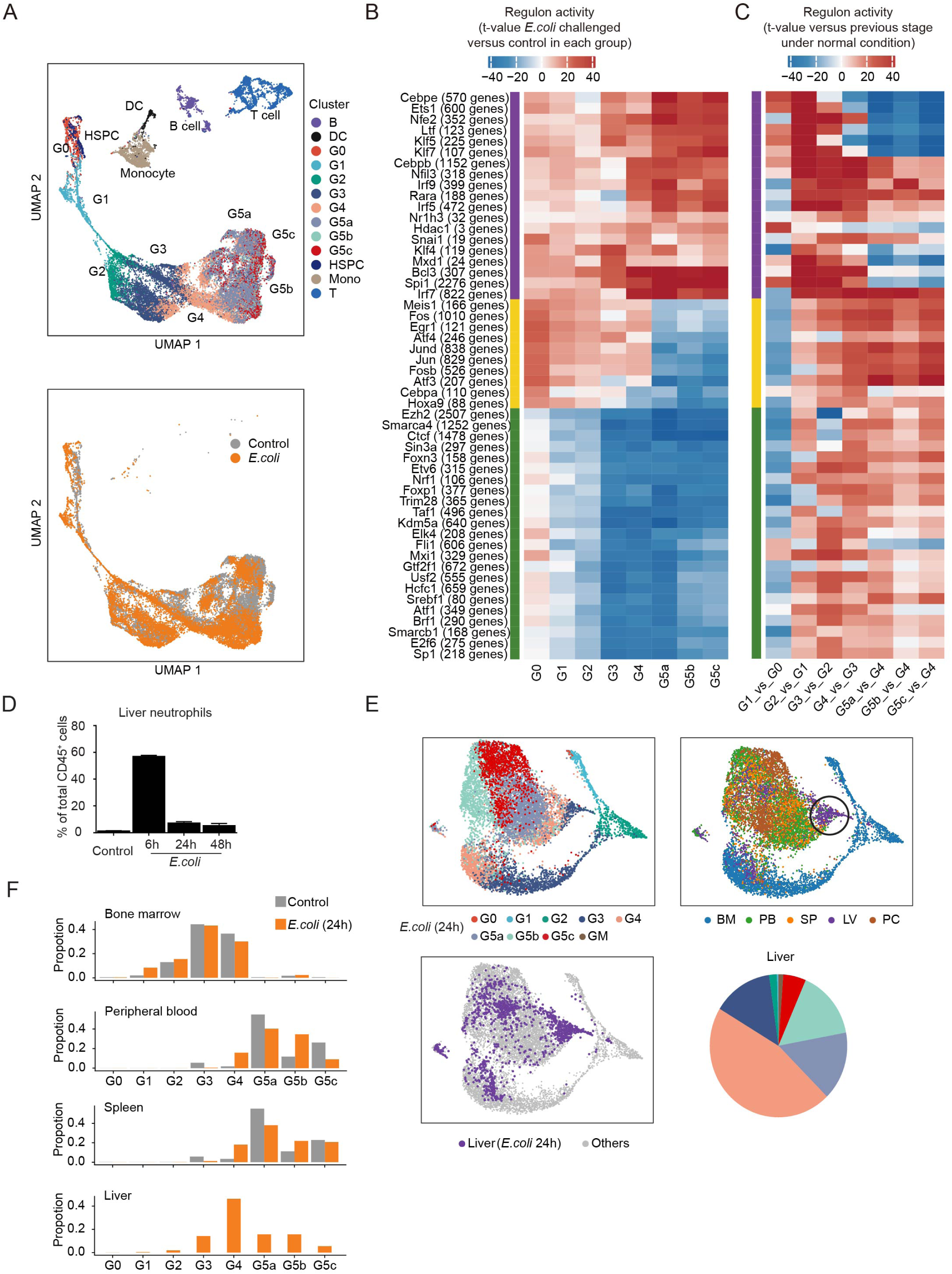
**(A-C) Alteration of transcription networks in *E. coli*-challenged mice.** (A) UMAP of the regulon activity matrix of 32,888 cells (11,992 normal neutrophils, 13,687 challenged neutrophils, and 7209 other cells under normal conditions) colored by Seurat cluster identity (left) or experimental condition (right, only neutrophils). (B) Heatmap of the t-values of regulon activity derived from a generalized linear model for the difference between cells from one challenged neutrophil subpopulation and cells from the corresponding normal subpopulation. Only regulons with at least one absolute t-value greater than 18 are visualized. Regulons are hierarchically clustered based on challenge-response pattern (purple: upregulated, yellow: first up-then downregulated, green: downregulated) (C) Heatmap showing activity change of regulons identified in (B) during normal group transitions. **(D-F) The liver displays a distinct extramedullary granulopoiesis program during bacterial infection.** (D) Neutrophil proportions in the liver measured at different time points after *E. coli* challenge. Results are the mean (±SD) of three independent experiments. (E) Composition of liver neutrophils after *E. coli* challenge. Top: UMAP of *E. coli*-challenged neutrophils from BM, PB, SP, liver (LV), and peritoneal cavity (PC) colored by cluster identity. Bottom: Liver cells are highlighted in the UMAP plot. Composition of the liver neutrophils after *E. coli* challenge is displayed on the right. (F) Comparisons of organ distribution of each neutrophil cluster before and after *E. coli* challenge.

**Figure S7.**
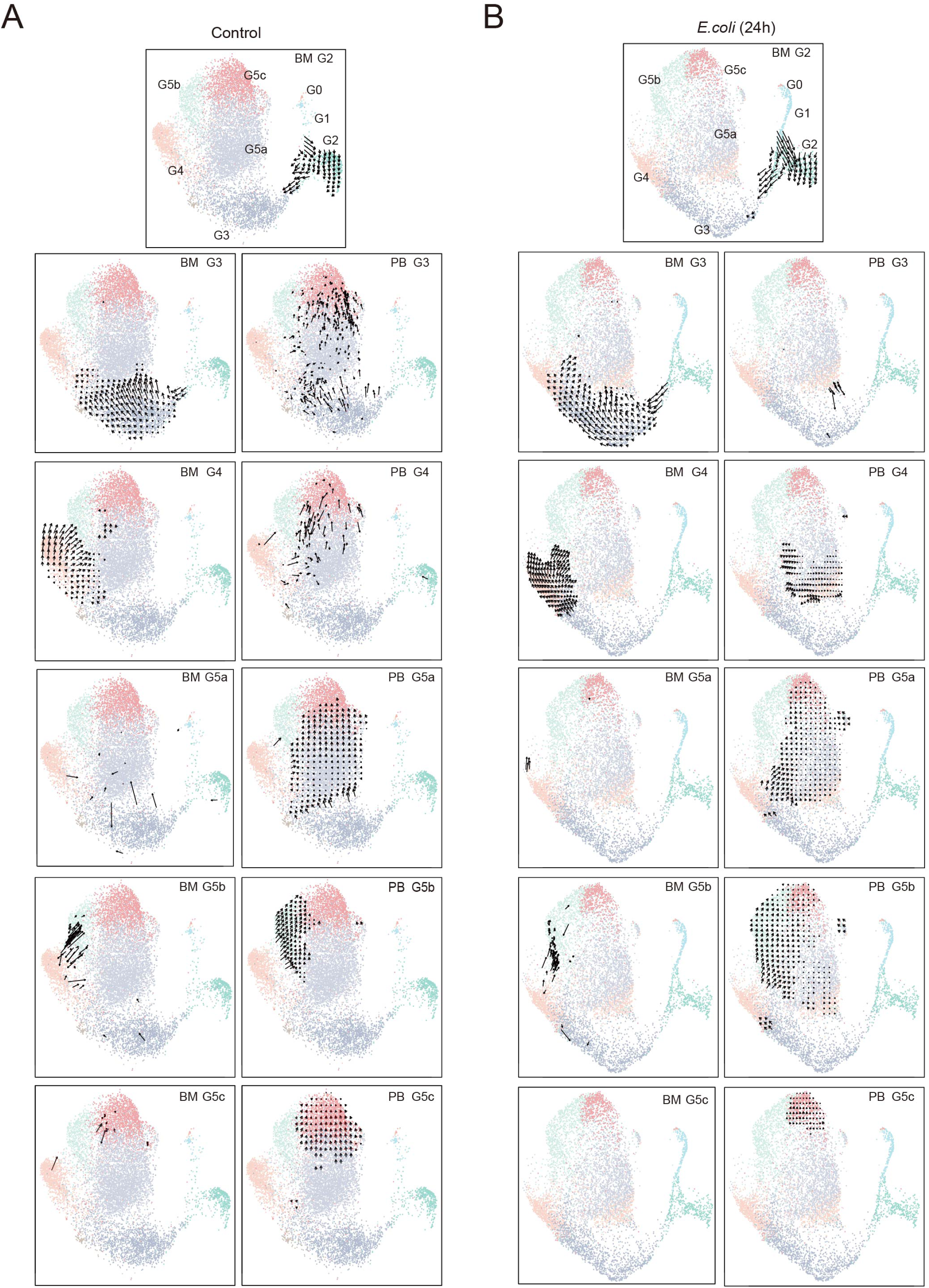
Neutrophil dynamics under steady-state and bacterial infection conditions assessed by velocity analysis. (A) Dynamics (velocity field projected on the UMAP plot) of G2-G5 neutrophils under normal conditions. BM neutrophils (left) and PB neutrophils (right) are displayed separately. For small populations, velocity vectors of all cells are visualized directly. For large populations, a grid velocity summary is derived by calculating the Gaussian-weighted average of velocity vectors of all cells at each grid point. Only grid points adjacent to neutrophils in the target population are visualized. (B) As in (A) but of challenged neutrophils.

